# Novel mechanism of inflammatory activation by Ebola virus matrix protein linked to the ebolavirus virulence

**DOI:** 10.1101/2024.10.06.616882

**Authors:** Satoko Yamaoka, Zeineb M’Hamdi, Lin Wang, Vaille A. Swenson, Kristin L. McNally, Shao-Chia Lu, Reema Singh, Stephanie L. Saundh, Brady N. Zell, Sonja M. Best, Michael A. Barry, Angela L. Rasmussen, Hideki Ebihara

## Abstract

Uncontrolled systemic inflammatory responses are a critical pathological feature of fatal Ebola virus (EBOV) infection. While some inflammatory responses may originate from mononuclear phagocytes (MNPs), non-immune cells vastly outnumber MNPs and may be an important source of inflammation. Here, we demonstrated that highly virulent EBOV induced a high and sustained pro-inflammatory response compared to less virulent ebolaviruses in non-MNPs through TLR4-independent NF-κB activation. We identified the EBOV matrix protein VP40 as a potent activator of NF-κB in non-MNPs, whose intrinsic inflammatory activation ability is higher than VP40 proteins from less virulent ebolaviruses. This suggests that VP40 is a novel virulence determinant inducing distinct degrees of pro-inflammatory responses among ebolaviruses. Mechanistically, VP40 activated the NF-κB signaling pathway, primarily via TNFR1 using a ligand-independent mechanism. These findings reveal mechanisms that may drive systemic inflammation and promote EBOV pathogenesis, suggesting potential therapeutic strategies to mitigate immune dysregulation in severe EBOV infections.

## Introduction

Ebola virus disease (EVD), caused by Ebola virus (EBOV), is one of the most severe viral diseases, with case fatality rates reaching as high as 90%^1^. The largest EVD outbreak in West Africa from 2013 to 2016 was devastating, with more than 28,600 people infected and more than 11,300 deaths^1,2^. Clinical studies indicate a strong correlation exists between uncontrolled pro-inflammatory activation and EVD fatality^3–11^. Uncontrolled pro-inflammatory responses associated with excessive production of various pro-inflammatory cytokines and chemokines, known as the cytokine storm, is a critical pathological feature of severe EVD^3,4,6–14^. This leads to systemic endothelial dysfunction, vascular leakage, and coagulation abnormalities, impairing the immune response against the viral infection^15–18^.

Several previous studies have demonstrated that immune cells, such as macrophages, dendritic cells (DCs), and T cells, play important roles in producing pro-inflammatory mediators during EBOV infection^13,19–21^. In mononuclear phagocytes (MNPs), inflammatory activation is initiated by EBOV surface glycoprotein (GP_1,2_) or by shed GP binding to toll-like receptor 4 (TLR4) ^21–34^, which activates the nuclear factor kappa B (NF-κB) pathway to drive pro-inflammatory responses^35,36^. Several studies have shown that EBOV GP binding to TLR4 triggers a temporary pro-inflammatory response in MNPs^13,24,25^ which may involve negative feedback loops that regulate an appropriate inflammatory response^28^.

TLR4 is only expressed on a small subset of host cells including MNPs^37^. Given this, inflammatory activation by GP through TLR4 may not explain the entire breadth and strength of uncontrolled inflammation during EBOV infection. This concept is supported by the observation that TLR4 inhibitors only partially reduce serum levels of some cytokines and chemokines in EBOV-infected mice^32^ and that TLR4 knockout mice are still fully susceptible to lethal EBOV infection^32^. Thus, EBOV-mediated induction of an uncontrolled pro-inflammatory response must involve as yet unidentified mechanisms in addition to the TLR4 engagement by GP.

In this study, we demonstrate that the EBOV matrix protein VP40 triggers sustained activation of NF-κB signaling via a tumor necrosis factor receptor (TNFR)-dependent mechanism in non-immune target cells lacking TLR4 expression. This finding represents a novel, TLR4-independent mechanism for amplifying and sustaining pro-inflammatory responses in the host by EBOV. Intriguingly, NF-κB activation induced by VP40 from EBOV is significantly higher than that elicited by VP40 from less virulent ebolaviruses such as Bundibugyo virus and Reston virus. This strongly suggests that VP40 may be a novel determinant of ebolavirus species-specific virulence. Our findings provide critical insights into the molecular mechanism leading to the induction of uncontrolled pro-inflammatory responses underlying EVD pathogenesis. Understanding these mechanisms is crucial for developing effective therapeutic strategies against severe EVD.

## Results

### EBOV infection induces significantly stronger pro-inflammatory responses compared to other ebolaviruses through a TLR4-independent mechanism

The genus *Orthoebolavirus* consists of six virus species, each represented by a single type virus: EBOV, Sudan virus (SUDV), Bundibugyo virus (BDBV), Taï Forest virus (TAFV), Reston virus (RESTV), and Bombali virus (BOMV)^2,38^. While EBOV and SUDV are highly virulent ebolaviruses, BDBV and TAFV are often referred to as moderately or less virulent ebolaviruses. RESTV is a unique ebolavirus thought not to cause clinically apparent disease in humans, although it can cause fatal disease in cynomolgus macaques^39^. The pathogenic potential of BOMV to humans remains undetermined (**Fig. 1a**).

**Fig. 1:**
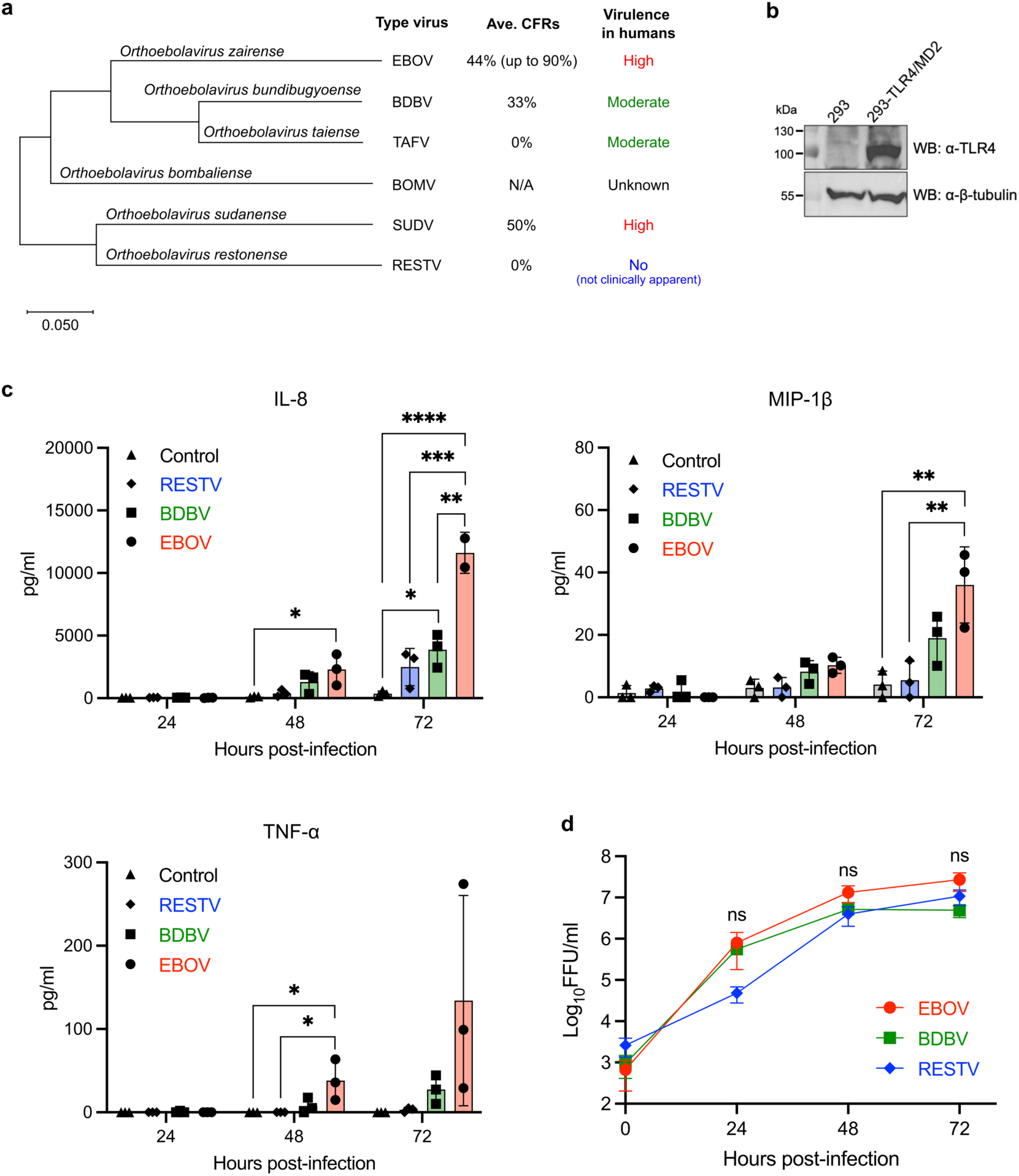
Species-specific differences in pro-inflammatory activation across ebolaviruses. **a** Phylogenetic tree of the genus *Orthoebolavirus* constructed with Mega X based on the nucleotide sequences of VP40 matrix protein. Average case fatality rates (Ave. CFRs) are calculated based on reported outbreaks as of April 2024. **b** Western blotting for TLR4 in 293 and 293-TLR4/MD2 cells. **c** Quantification of IL-8, MIP-1β, and TNF-α released in the supernatants of EBOV, BDBV, or RESTV-infected 293 cells (MOI = 1). **d** Viral growth kinetics in 293 cells (MOI = 1). The Virus infectivity titers in culture supernatants are shown in Log_10_ focus-forming unit (FFU)/ml. Control: Mock-infected. For **c** and **d**, data are shown with mean ± SD (n = 3 independent biological replicates). ns > 0.05, \**P* ≤ 0.05, \*\**P* ≤ 0.01, \*\*\**P* ≤ 0.001, \*\*\*\**P* ≤ 0.0001; ordinary one-way ANOVA for each time point.

We first compared the ability of EBOV, BDBV, and RESTV to induce a pro-inflammatory response in 293 cells which do not express detectable levels of TLR4^27,40^ (**Fig. 1b**) and have been suggested to originate from the adrenal glands^41^, one of the main target organs for EBOV. EBOV infection induced a time-dependent increase in the release of pro-inflammatory chemokines IL-8 and MIP-1β, as well as pro-inflammatory cytokine TNF-α (**Fig. 1c**). Notably, the levels of IL-8, MIP-1β, and TNF-α were consistently higher in cells infected with EBOV when compared to those induced by BDBV and RESTV (**Fig. 1c**). No significant difference in virus replication was observed among the three ebolaviruses (**Fig. 1d**). These results indicate that highly virulent EBOV triggers the pro-inflammatory activation to a greater extent than the less virulent BDBV or non-pathogenic RESTV through a mechanism independent of TLR4.

### NF-κB is critically involved in the varying degrees of pro-inflammatory activation observed across ebolaviruses

We next performed RNA sequencing (RNA-seq) to investigate global transcriptomic changes in 293 cells infected with EBOV and RESTV, ebolaviruses with distinct pathogenicity in humans. A multi-dimensional scaling (MDS) plot revealed wide separation in gene expression profiles between EBOV and RESTV infections at 48 hours post-infection (hpi), indicating substantially different global expression profiles in terms of directionality and magnitude of expression (**Fig. 2a**). Indeed, a smaller number of both upregulated and downregulated differentially expressed genes (DEGs) were observed in RESTV infection compared to EBOV infection at 48 hpi when both were compared to uninfected cells (**Fig. 2b**, **Supplementary Data 1**). Gene set enrichment analysis (GSEA) utilizing the Hallmark gene sets demonstrated pathways related to inflammation, such as TNFα Signaling via NFκB, were enriched in both EBOV and RESTV infections, however a greater number of genes were enriched in response to EBOV infection (**Fig. 2c**, **Supplementary Fig. 1a**). Functional analysis with Ingenuity Pathway Analysis (IPA) predicted numerous inflammatory signaling pathways, such as IL-6, IL-1, TNFR1, TNFR2 Signaling, and NFκB Activation by Viruses, to be more strongly activated in EBOV than in RESTV (**Fig. 2d**, **Supplementary Data 2**). In addition, the IPA Upstream Analysis, supported by subsequent DE analyses, indicated that NF-κB-driven inflammatory gene expression was more enriched upon EBOV infection than RESTV (**Fig. 2e**, **Supplementary Fig. 1b-c, 2a-b**).

**Fig. 2:**
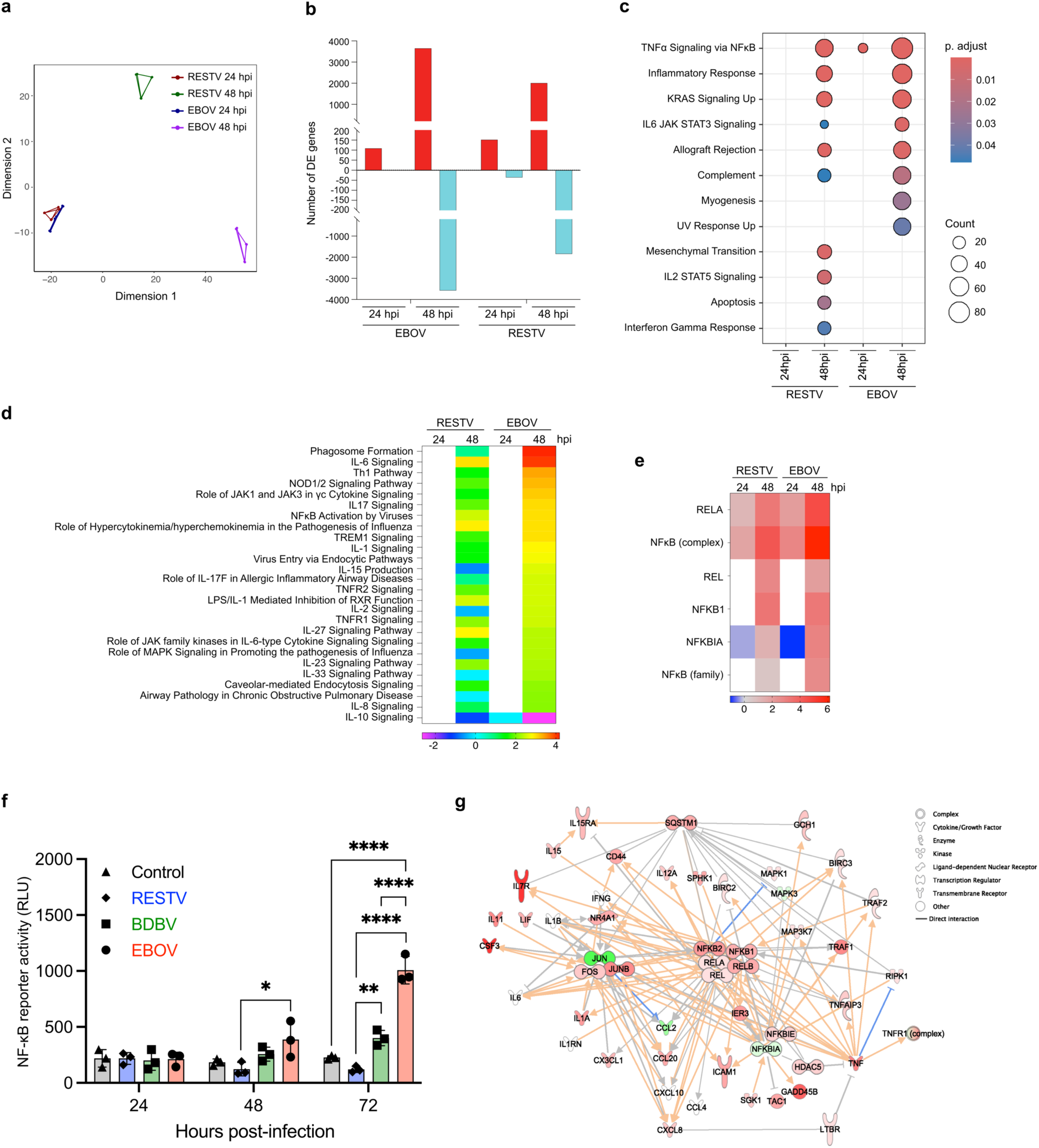
Distinct degree of NF-κB activation induced by EBOV compared to other ebolaviruses. **a** MDS plot showing DEGs in RESTV-and EBOV-infected 293 cells. MDS plot depicts two sets of three replicates corresponding to samples run in different sequencing lanes. **b** Number of genes meeting DE criteria (fold change relative to time-matched mock-infected controls > |1.5|, adjusted *P*-value < 0.05). Red represents upregulated and light green represents downregulated DEGs. **c** Dot plot for upregulated DEGs generated based on GSEA using hallmark gene sets. Gene sets were sorted to only include genes that reached DE criteria within the EBOV 48 hpi dataset. **d** Ingenuity Pathway Analysis (IPA) Canonical Pathway Analysis showing Z-scores for predicted activation state of significantly enriched pathways (*P* < 0.01). Heatmap shows pathways with z > |2| from the “Cytokine Signaling” and “Pathogen-Influenced Signaling” subcategories of IPA Signaling Pathways. Red and blue indicate pathway activation and inhibition. White indicates the pathway did not meet enrichment criteria at this time point. The unfiltered pathway list is in **Supplementary Data 2**. **e** IPA Upstream Analysis showing Z-scores for predicted upstream activation state of NF-κB family members (*P* < 0.01). Heatmap shows transcriptional regulators with z > |2| from the NF-κB family. Red and blue indicate predicted activation and inhibition of downstream targets. White indicates that the upstream regulator did not meet downstream enrichment criteria at this time point. The unfiltered list of transcriptional regulators is in **Supplementary Data 3**. **f** NF-κB-responsive luciferase reporter activity in EBOV-, BDBV-, or RESTV-infected 293 cells. Control: Mock-infected. **g** IPA network built from NF-κB-associated genes at 48 hpi for EBOV. Red and green indicate positive (upregulation) and negative fold change (downregulation) relative to the time-matched mock-infected controls. Darker shading indicates genes met DE criteria, lighter shading indicates they did not. No shading indicates no change relative to controls. Orange and blue lines indicate predicted activation and inhibition. Gray lines indicate insufficient information to predict molecular activity. Solid lines indicate known direct interactions. Arrowheads indicate directionality. For **f**, data are shown with mean ± SD (n = 3 independent biological replicates). \**P* ≤ 0.05, \*\**P* ≤ 0.01, \*\*\*\**P* ≤ 0.0001; ordinary one-way ANOVA for each time point.

Given the established crucial role of NF-κB in driving pro-inflammatory activation^35,36^, we proceeded to experimentally verify NF-κB activation induced by ebolavirus infection. A NF-κB-responsive luciferase reporter assay demonstrated a time-dependent increase in NF-κB activity induced by EBOV, significantly higher than that induced by BDBV and RESTV at 72 hpi (**Fig. 2f**). While BDBV infection induced a modest increase in NF-κB activity over time, RESTV infection did not elicit any discernible increase in NF-κB activity. The central role of NF-κB in pro-inflammatory responses induced by EBOV was further visually demonstrated using a network assembled from transcriptome IPA results, focusing on the ‘NFκB Activation by Viruses’ pathway (**Fig. 2g**). TLR4-related pathways did not meet enrichment or activation criteria (**Supplementary Data 2**), verifying a lack of expression in the 293 cells used in this analysis. Together, these results suggest that NF-κB is the key regulator for driving different degree of pro-inflammatory responses between EBOV and other ebolaviruses.

### EBOV matrix protein VP40 activates NF-κB and induces the production of pro-inflammatory mediators in non-MNP target cells

To examine which EBOV protein(s) might mediate NF-κB activation, we individually expressed seven structural EBOV proteins (NP, VP35, VP40, GP, VP30, VP24, and L) in 293 cells and assessed NF-κB activation ability of each viral protein. Among the seven viral proteins, the matrix protein VP40 was the only EBOV protein that activated NF-κB (**Fig. 3a**, **Supplementary Fig. 3**). VP40-mediated NF-κB activation was also verified in two other human cell lines, Huh7 and HepG2 hepatocytes, which also represent main target cell type for EBOV infection^42–46^ (**Fig. 3b**). Notably, while the magnitude of NF-κB activation was similar following expression of VP40 and TRAF6 used as a key signal transducer in the NF-κB pathway^47^, the NF-κB activation induced by VP40 remained elevated for at least 72 hours post-transfection (hpt), whereas TRAF6 induced transient NF-κB activation that peaked by 24 hpt and subsequently declined (**Fig. 3c**).

**Fig. 3:**
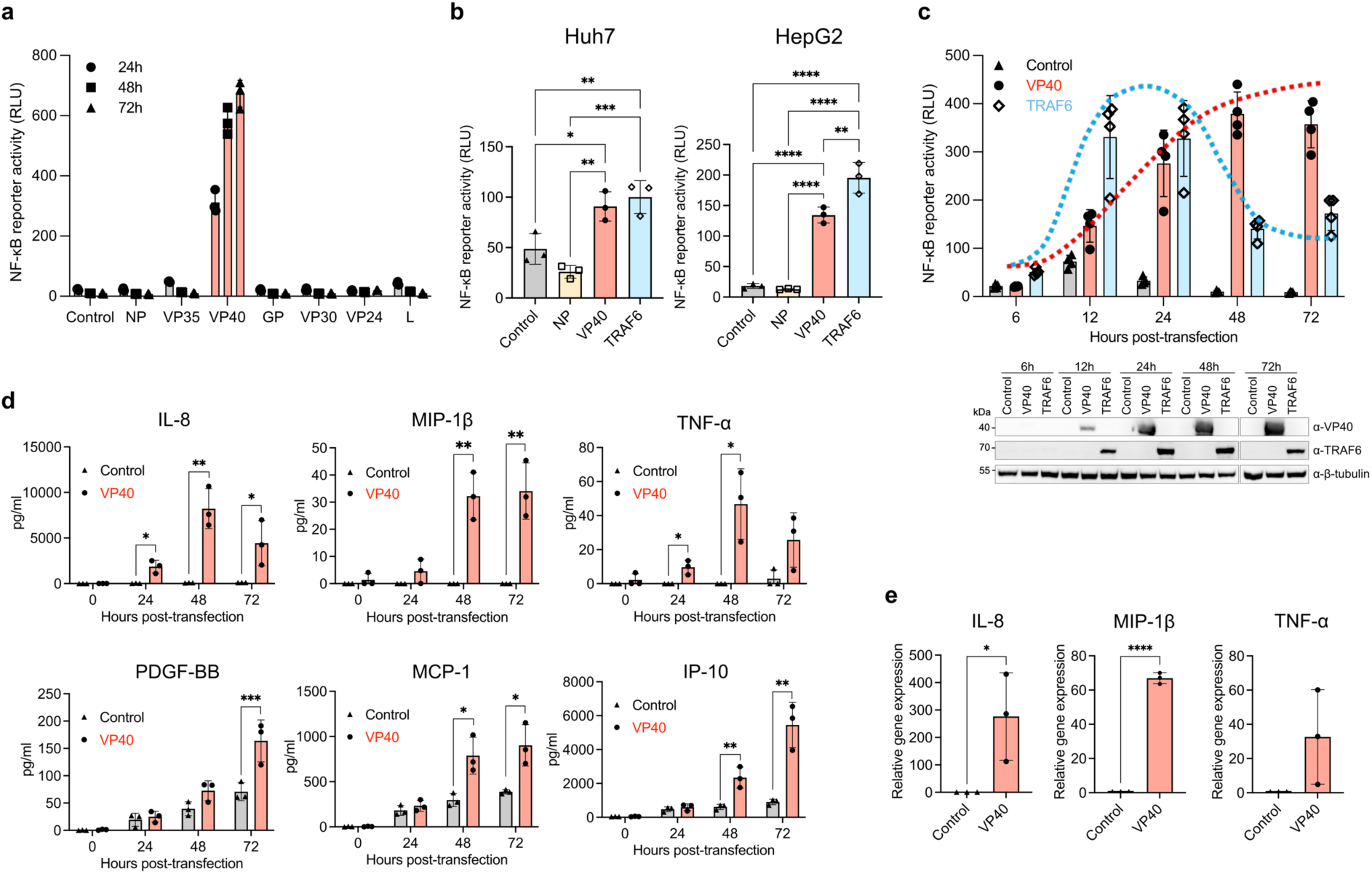
Pro-inflammatory and NF-κB activation elicited by EBOV VP40 individual expression. **a** NF-κB-responsive luciferase reporter activity in EBOV protein-expressing 293 cells. **b** NF-κB-responsive luciferase reporter activity in EBOV NP- or EBOV VP40-, or TRAF6-expressing Huh7 cells (left) and HepG2 cells (right) at 24 hpt. **c** Time course of NF-κB-responsive luciferase reporter activity in EBOV VP40- or TRAF6-expressing 293 cells. **d** Quantification of IL-8, MIP-1β, TNF-α, PDGF-BB, MCP-1, and IP-10 released in the supernatants of EBOV VP40-expressing 293 cells. **e** Quantification of IL-8, MIP-1β, and TNF-α mRNA in EBOV VP40-expressing 293 cells at 48 hpt. Control: Empty vector transfected. Data are shown with mean ± SD (n = 3 independent biological replicates). \**P* ≤ 0.05, \*\**P* ≤ 0.01, \*\*\**P* ≤ 0.001, \*\*\*\**P* ≤ 0.0001; ordinary one-way ANOVA for **b** and unpaired t test for each time point for **d** and **e**.

Importantly, VP40-expressing 293 cells demonstrated the release of pro-inflammatory mediators in a pattern generally resembled to that observed in EBOV-infected cells (**Fig. 3d**); the expression levels of IL-8, MIP-1β, and TNF-α were all significantly higher in cells expressing VP40 compared to the control group transfected with an empty vector, while PDGF-BB, MCP-1, and IP-10 were also increased. We quantified the mRNA levels of key inflammatory mediators, such as IL-8, MIP-1β, and TNF-α, and found significant upregulation in cells expressing VP40 compared to the control group (**Fig. 3e**). These findings highlight the significant role of VP40 in pro-inflammatory activation through NF-κB in non-MNP target cells.

### The distinct inflammatory phenotypes of ebolaviruses correlate with their respective abilities of VP40 for NF-κB activation

We next examined whether VP40 proteins from different ebolaviruses activate NF-κB to different degrees. Notably, NF-κB reporter activity induced by EBOV VP40 was significantly higher than that elicited by VP40 from other ebolaviruses (**Fig. 4a**). While EBOV VP40 induced dose-dependent NF-κB activation, such dose-dependency was not observed for VP40 from other ebolaviruses, particularly for RESTV VP40. Even at the maximum tested dose (2 μg VP40-expression plasmid), NF-κB reporter activity induced by RESTV VP40 did not reach the level achieved by EBOV VP40 (**Fig. 4b**). The robust ability of VP40 to activate NF-κB was conserved across several EBOV variants, including Mayinga, Kikwit, and Makona-C07 (**Supplementary Fig. 4**). These results indicate that there is an association between virulence/inflammatory phenotypes of the ebolaviruses and their respective VP40 for NF-κB activation.

**Fig. 4:**
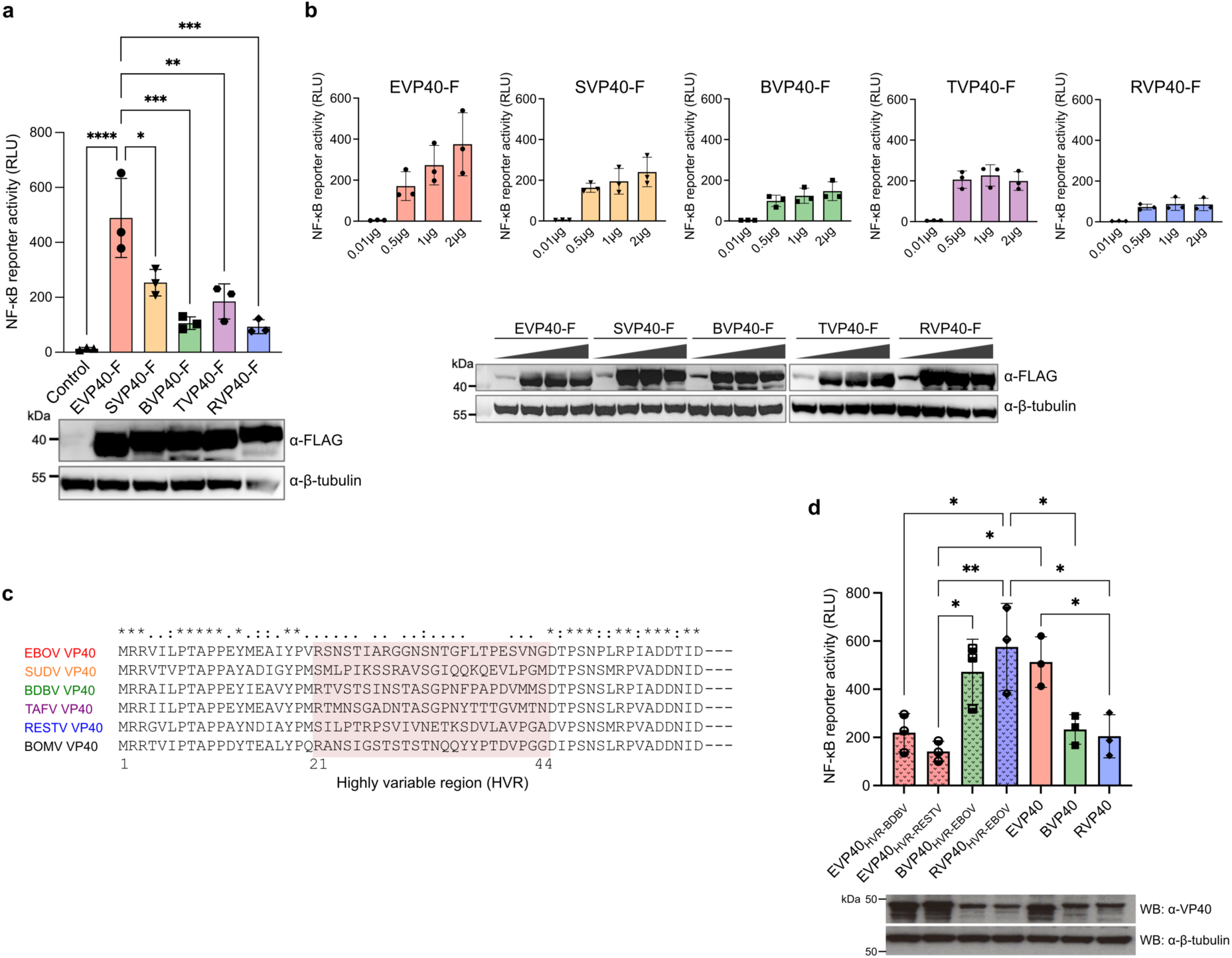
Differing ability in NF-κB activation among VP40 from different ebolaviruses. NF-κB-responsive luciferase reporter activity in 293 cells expressing FLAG-tagged VP40 derived from EBOV, SUDV, BDBV, TAFV, or RESTV **a** at 72 hpt, and **b** at 48 hpt for the plasmid titration. **c** Amino acid sequence alignment of VP40 from EBOV, SUDV, BDBV, TAFV, RESTV, and BOMV generated using SnapGene. The sequence conservation is indicated by symbols. **d** NF-κB-responsive luciferase reporter activity in 293 cells expressing either the parental VP40 derived from EBOV, BDBV, RESTV, or chimeric VP40 proteins where the HVR of EBOV VP40 was replaced with that of BDBV or RESTV (EVP40_HVR-BDBV_ and EVP40_HVR-RESTV_), as well as chimeric BDBV and RESTV VP40 proteins containing the HVR of EBOV (BVP40_HVR-EBOV_ and RVP40_HVR-EBOV_) at 72 hpt. A polyclonal antibody targeting the EBOV VP40 was employed as the primary antibody in the Western blotting analysis. Control: Empty vector transfected. Data are shown with mean ± SD (n = 3 independent biological replicates). \**P* ≤ 0.05, \*\**P* ≤ 0.01, \*\*\**P* ≤ 0.001, \*\*\*\**P* ≤ 0.0001; ordinary one-way ANOVA.

### The N-terminal highly variable region of VP40 determines its ability to activate NF-κB

Alignment of the amino acid sequences of VP40 from all six ebolaviruses revealed a high sequence similarity across the protein (**Supplementary Fig. 5**), with the exception of a highly variable region (HVR) located at the N-terminus, amino acid (aa) positions 21 to 44 (**Fig. 4c**). Intriguingly, EBOV VP40 chimeras possessing the HVR of BDBV or RESTV VP40 (EVP40_HVR-BDBV_ and EVP40_HVR-RESTV_) showed a reduced ability to activate NF-κB, to a degree similar to wild-type BDBV and RESTV VP40 (**Fig. 4d**). Likewise, BDBV and RESTV VP40 chimeras possessing the EBOV HVR (BVP40_HVR-EBOV_ and RVP40_HVR-EBOV_) activated NF-κB to a degree comparable to wild-type EBOV VP40 (**Fig. 4d**). The mutant EBOV VP40, VP40_Δ1-20aa_, which lacks the first 20 aa residues at the N-terminus, including the two classical late domain motifs, 7-PTAP-10 and 10-PPEY-13, retained its ability to activate NF-κB (**Supplementary Fig. 6a-b**). These findings indicate the critical role of the 24 aa HVR within EBOV VP40 in activating NF-κB, while the two classical late domains are dispensable.

### EBOV VP40 activates the p65-dependent canonical NF-κB pathway

There are two primary signaling pathways, both canonical and non-canonical, which are known to activate NF-κB^35,36^ (**Fig. 5a**). We initially investigated which NF-κB subunit(s) translocate to the nucleus upon EBOV VP40 expression (**Fig. 5b**). Over-expression of positive control TRAF6 led to the nuclear localization of all tested NF-κB subunits, including p65, RelB, cRel, p52, and p50, verifying the involvement of both canonical and non-canonical pathways for NF-κB activation^47^. In contrast, in cells expressing VP40, p65, cRel, and p50 were observed to translocate to the nucleus, while RelB and p52 did not show such localization. This suggests that VP40-mediated NF-κB activation is driven by the canonical pathway, but not the non-canonical pathway. Furthermore, a significant increase in p52 expression levels was observed with TRAF6 or NIK over-expression, but not with VP40 (**Supplementary Fig. 7a**), thereby ruling out the possibility of VP40 activating the non-canonical NF-κB pathway via the NIK-IKKα axis.

**Fig. 5:**
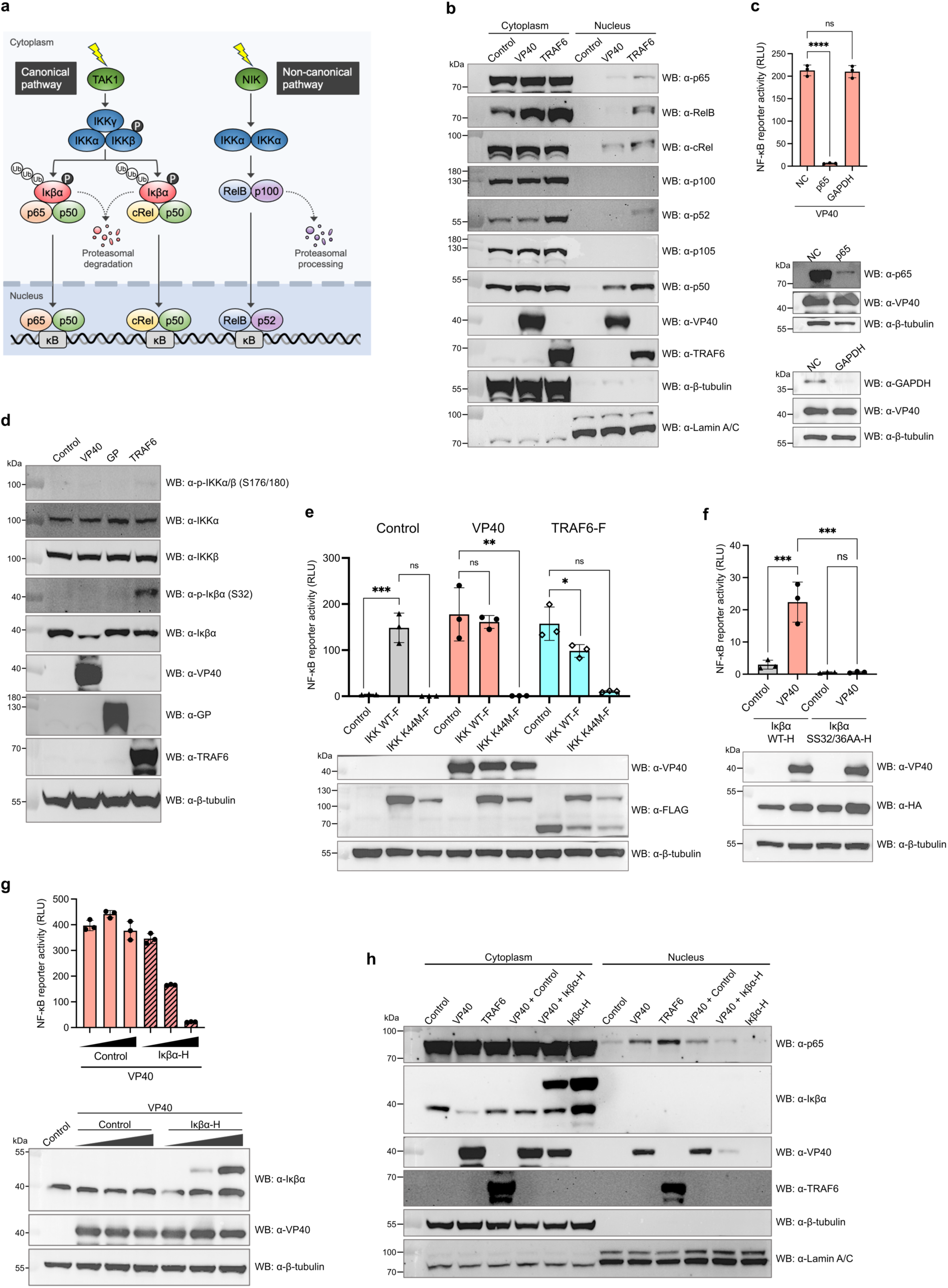
Examination of IKK-p65 axis activation in the canonical NF-κB signaling pathway in the presence of EBOV VP40. **a** Diagram of the canonical and non-canonical NF-κB pathways. The canonical pathway involves the IκB kinase (IKK) complex, which comprises IKKα, IKKβ, and IKKγ. Activation of the IKK complex leads to the phosphorylation/ubiquitination and subsequent proteasomal degradation of Iκβα, allowing either p65:p50 or cRel:p50 NF-κB heterodimers to enter the nucleus. The non-canonical pathway relies on the cooperation of NF-κB-inducing kinase (NIK) and its downstream kinase IKKα. Activation of NIK and IKKα triggers the proteasomal processing of p100 into p52, leading to the nuclear translocation of RelB:p52 NF-κB heterodimers. **b** Western blotting for NF-κB subunits in cytoplasmic and nuclear fractions isolated from 293 cells expressing EBOV VP40 or TRAF6 at 24 hpt. **c** NF-κB-responsive luciferase reporter activity in 293 cells expressing EBOV VP40 at 48 hpt, following siRNA-mediated knockdown of p65 or GAPDH. **d** Western blotting for IKK complex and Iκβα in 293 cells expressing EBOV VP40, EBOV GP, or TRAF6 at 24 hpt. **e** NF-κB-responsive luciferase reporter activity in 293 cells expressing 0.25 μg of IKK wild-type or IKK mutant (K44M) together with either 0.25 μg of EBOV VP40 or TRAF6 at 48 hpt. **f** NF-κB-responsive luciferase reporter activity in Iκβα-knockout 293 cells expressing 0.25 μg of Iκβα wild-type or Iκβα mutant (SS32/36AA) together with 0.25 μg of EBOV VP40 at 48 hpt. **g** NF-κB-responsive luciferase reporter activity in 293 cells expressing 0.01, 0.1, or 0.5 μg of Iκβα together with 0.5 μg of EBOV VP40 at 48 hpt. **h** Western blotting for p65 in cytoplasmic and nuclear fractions isolated from 293 cells expressing EBOV VP40 with or without Iκβα, TRAF6, or Iκβα at 24 hpt. Control: Empty vector transfected. For **c** and **e**-**g**, data are shown with mean ± SD (n = 3 independent biological replicates). ns > 0.05, \**P* ≤ 0.05, \*\**P* ≤ 0.01, \*\*\**P* ≤ 0.001, \*\*\*\**P* ≤ 0.0001; ordinary one-way ANOVA.

The p65:p50 heterodimer, with p65 acting as a key transcription factor, is recognized as the primary form of NF-κB that activates pro-inflammatory genes via the canonical pathway^49^. We examined whether the p65 NF-κB subunit is necessary for NF-κB activation induced by VP40. Depleting p65 through siRNA transfection in 293 cells markedly reduced NF-κB activity in cells expressing VP40 (**Fig. 5c**). Taken together, these findings indicate that VP40 activates the p65-dependent canonical NF-κB pathway.

### IKK activation is essential for NF-κB signaling triggered by EBOV VP40

We next investigated the upstream signaling events in the canonical NF-κB pathway, including IKK activation followed by Iκβα degradation (**Fig. 5a**). The expression level of Iκβα was significantly reduced in the presence of VP40 at 24 hpt compared to the negative controls, such as empty vector or EBOV GP expression (**Fig. 5d**). The time course analysis further revealed that the Iκβα reduction became detectable starting from 18 hpt, becoming more pronounced by 24 hpt (**Supplementary Fig. 7b**), despite an increase in Iκβα mRNA levels over time (**Supplementary Fig. 7c**). In contrast, the reduction of Iκβα level was not notable upon expression of RESTV VP40 (**Supplementary Fig. 7d**), nor in the expression of EBOV VP40 chimeras, such as EVP40_BDBV-HVR_ and EVP40_RESTV-HVR_ (**Supplementary Fig. 7e**). The expression of EVP40_Δ1-20_ reduced Iκβα levels to a similar level as wild-type EBOV VP40 (**Supplementary Fig. 7e**). Although western blotting did not clearly detect phosphorylation of Iκβα and IKKα/β in the cells expressing EBOV VP40, two supplementary experiments revealed the critical roles of Iκβα phosphorylation and IKKα/β kinase activity in EBOV VP40-mediated NF-κB activation. First, the expression of a kinase-inactive, dominant negative IKKβ mutant (K44M) significantly impaired the NF-κB activity induced by VP40 as well as TRAF6 (**Fig. 5e**). Second, replacing the endogenous, wild-type Iκβα by overexpression of a canonical phosphorylation site-deficient Iκβα mutant (SS32/36AA) in Iκβα CRISPR-Cas9 knockout cells resulted in a complete inhibition of VP40-mediated NF-κB activation (**Fig. 5f**). These findings strongly suggest that VP40-induced NF-κB activation is mediated by IKKα/β kinase and Iκβα phosphorylation in the canonical NF-κB pathway. The absence of phosphorylation on Iκβα and IKKα/β on our western blotting results may be attributed to their expression levels, which could result from their rapid degradation, or some technical factors, such as limitations in detection sensitivity.

The involvement of Iκβα reduction in NF-κB activation was further validated by an observed negative correlation between Iκβα levels and NF-κB reporter activity in the VP40-expressing cells (**Fig. 5g**). Moreover, cytoplasmic and nuclear fractionation assay demonstrated that co-expression of Iκβα with VP40 resulted in a reduction of p65 nuclear accumulation compared to VP40 expression alone (**Fig. 5h**). Together, these results suggest that VP40 triggers IKK activation followed by Iκβα phosphorylation and degradation, leading to the pro-inflammatory gene expression regulated by NF-κB.

### TNFR1 contributes to the NF-κB activation induced by EBOV VP40

Multiple cellular receptors, such as TNFR superfamily members (TNFRSF)^50^, interleukin receptors (ILRs)^51^, and NOD-like receptors (NLRs)^52^ initiate signal transduction cascades after ligand binding to activate the IKK-p65 axis in the canonical NF-κB signaling pathway. IPA Upstream Analysis filtered by cytokines indicated the involvement of TNF in inflammatory cascade activation during EBOV infection (**Fig. 6a**). In addition, TNF was predicted to be a much stronger upstream regulator in EBOV infection when compared to RESTV (**Fig. 6a**). The IPA analyses also indicated a pronounced role of TNFR1 in NF-κB activation after EBOV infection (**Supplementary Fig. 8**). Together, these findings indicate that TNF signaling pathway is activated during EBOV infection.

**Fig. 6:**
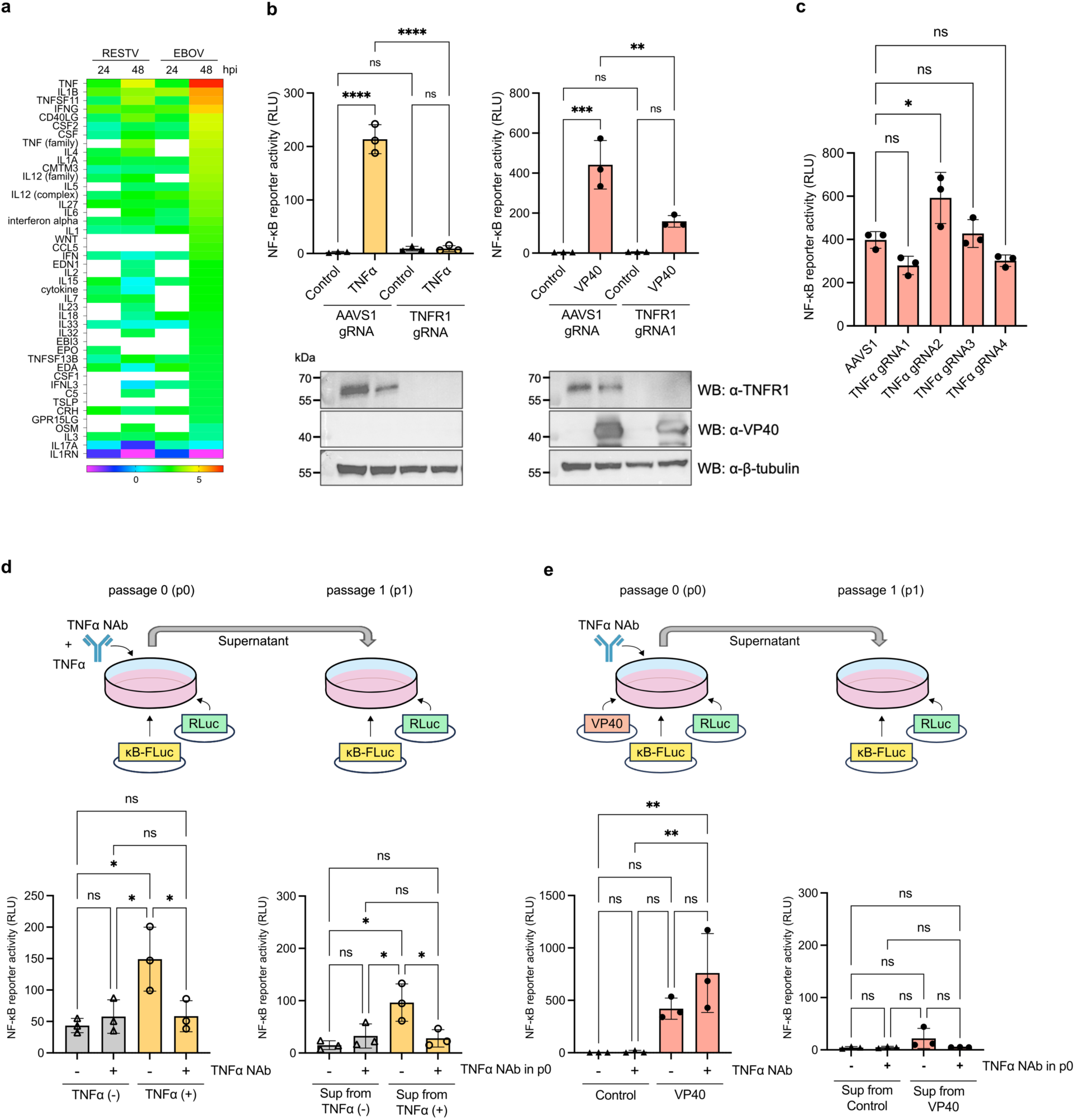
Identification of upstream regulator involving NF-κB signaling activation induced by EBOV VP40. **a** IPA Upstream Analysis sorted by cytokines. Z-scores indicating predicted activity of cytokines upstream of significantly enriched DEGs in the dataset (enrichment significance threshold set at *P* < 0.01). Heatmap shows cytokines with z > |2| and is shaded across a rainbow spectrum with red and orange indicating activation and blue and purple indicating inhibition. White indicates that the pathway did not meet enrichment or z-score criteria at this time point. **b** NF-κB-responsive luciferase reporter activity in TNFR1-knockout 293 cells treated with 15 ng/ml of TNFα for 6 hours (left) and cells expressing EBOV VP40 at 48 hpt (right). **c** NF-κB-responsive luciferase reporter activity in TNFα-knockout 293 cells expressing EBOV VP40 at 48 hpt. **d** NF-κB-responsive reporter activity in 293 cells treated with mixture of TNFα and TNFα NAb for 6 hours (left; p0 cells) and cells treated with supernatant from p0 cells for 8 hours (right; p1 cells). **e** NF-κB-responsive reporter activity in EBOV VP40-expressing 293 cells treated with TNFα NAb at 48 hpt (left; p0 cells) and cells treated with supernatant from p0 cells for 8 hours (right; p1 cells). Control: Empty vector transfected. For **b**-**e**, data are shown with mean ± SD (n = 3 independent biological replicates). ns > 0.05, \**P* ≤ 0.05, \*\**P* ≤ 0.01, \*\*\**P* ≤ 0.001, \*\*\*\**P* ≤ 0.0001; ordinary one-way ANOVA.

To examine whether TNFR1 is involved in the NF-κB activation induced by EBOV VP40, we generated TNFR1 knockout cells using CRISPR-Cas9 gene editing and performed an NF-κB reporter assays. Notably, VP40-induced NF-κB activity was significantly reduced in TNFR1 knockout cells compared to a negative control cell line that was edited with CRISPR-Cas9 targeting the Adeno-Associated Virus Integration Site 1 (AAVS1) (**Fig. 6b**; **right, Supplementary Fig. 9a, c**). Despite a significant reduction, TNFR1 knockout did not completely abolish VP40-mediated NF-κB activation, as seen in TNFα-mediated NF-κB activation (**Fig. 6b**; **left**). Based on this result, we also knocked out another TNFRSF member, lymphotoxin β receptor (LTβR), which was reported to be expressed in 293 cells^50^. This additional knockout of LTβR further reduced the NF-κB activity induced by VP40 (**Supplementary Fig. 9b-c**). An IPA custom network with genes associated with VP40-mediated NF-κB activation reaffirmed that TNFR1, LTβR, NF-κB and their interacting proteins, including inflammatory cytokines, such as TNF, are upregulated after EBOV infection compared to RESTV (**Supplementary Fig. 9d**). These findings strongly suggest that several TNFRSF members additively contribute to the VP40-mediated NF-κB activation in EBOV-infected cells.

### VP40-mediated NF-κB activation by TNFR1 is independent of its ligand TNFα

Given that TNFR1 activation is typically triggered by binding of its ligand TNFα^53^, we subsequently tested the contribution of TNFα in the VP40-mediated NF-κB activation. Four clones of TNFα CRISPR-Cas9 knockout cell lines were established and the knockout effect was confirmed by flow cytometry staining and genomic DNA sequencing (**Supplementary Fig. 10, 11d**). There was no statistically significant reduction in VP40-mediated NF-κB activation in these TNFα knockout cell lines (**Fig. 6c**). We next examined whether TNFα secreted from VP40-expressing cells contributes additively to the activation of NF-κB. To test this, cells were treated with TNFα neutralizing antibody (NAb) (passage 0; p0 cells), and then their cell supernatants were used to stimulate TNFR1 on the newly seeded cells (passage 1; p1 cells). Positive control TNFα induced NF-κB activation, which was markedly reduced in both p0 and p1 cells when TNFα was combined with TNFα NAb (**Fig. 6d**). In contrast, TNFα NAb had no negative effect on VP40-mediated NF-κB activation in p0 cells (**Fig. 6e**). Additionally, NF-κB activity in p1 cells, which received supernatants from VP40-expressing p0 cells, was significantly lower compared to that in the VP40-expressing p0 cells (**Fig. 6e**). In conclusion, these findings demonstrate that, despite the significant contribution of TNFR1, VP40-mediated NF-κB activation is not triggered by autocrine and paracrine TNFα signaling through TNFR1. This strongly suggests that EBOV VP40 activates the NF-κB pathway primarily through TNFR1 but with a ligand-independent mechanism.

## Discussion

Here, we report a novel mechanism for the activation of a sustained pro-inflammatory response by EBOV. Systemic, sustained, and dysregulated inflammatory responses induced by EBOV infection are critical drivers of fatal disease progression in EVD^3,4,6–14^. Importantly, our study demonstrates that the EBOV matrix protein VP40 induces sustained pro-inflammatory responses by activating the canonical NF-κB signaling pathway, primarily via TNFR1 with a TNFα ligand-independent mechanism. Of note, the ability of EBOV VP40 to activate pro-inflammatory responses was found in several non-MNP target cells, including cells derived from human adrenal glands^41^ and hepatocytes. While monocytes and dendritic cells are critical as the initial target cells for EBOV infection^15,42,54,55^, subsequent migration of virus to target organs (e.g. liver, lymph nodes, spleen, adrenal gland) results in robust viral replication and systemic immunopathology^42–46^. Organ-associated inflammatory dysregulation has been observed in various lethal EVD models^45,56,57^, suggesting the significance of non-MNP target cells in amplifying systemic inflammatory responses in the late phase of the disease. Moreover, TNFR1, a receptor involved in VP40-mediated NF-κB activation, is a ubiquitous membrane receptor, and its expression can be found in various cell types *in vivo*^53^. This contrasts with TLR4 − a receptor involved in GP-mediated NF-κB activation − which is mainly expressed in MNPs^37^. Our study suggests that the coordinated mechanism of EBOV GP-mediated activation of inflammatory responses in MNPs and VP40-mediated amplification of inflammatory responses in non-MNP cells could be a driving factor in the development of a cytokine storm during EBOV infection *in vivo*.

The present study also proposes that VP40 is a potential novel virulence factor in determining distinct degrees of pro-inflammatory responses among ebolaviruses. Several clinical and experimental studies have suggested distinct patterns of inflammatory activation induced by each ebolavirus with different pathogenic potentials in humans. For instance, the expression levels of pro-inflammatory mediators in patients infected with SUDV or BDBV are shown to be lower than those in fatalities infected by EBOV^58,59^. Moreover, a significantly weaker inflammatory response was observed in BDBV-or RESTV-infected human peripheral blood mononuclear cells, compared to EBOV infection^29,60,61^. A potential role for GP in virus-specific induction of inflammation was suggested by Olejnik et al., demonstrating that, unlike EBOV GP, RESTV GP does not trigger TLR4 signaling in primary human monocyte-derived macrophages^29^. This finding suggests that GP-mediated TLR4 activation might be a critical factor in determining the different inflammatory activations induced by EBOV or RESTV in macrophages. However, the finding described herein that VP40 derived from the most virulent ebolavirus, EBOV, exhibits a greater ability to activate pro-inflammatory responses via NF-κB signaling than VP40 derived from other less virulent ebolaviruses, such as RESTV and BDBV, in non-immune cells. This may provide a mechanism for systemic inflammation and strongly suggests that VP40 serves as a novel key determinant for species-specific differences in inflammatory activation and virulence among ebolaviruses in humans.

To date, only limited studies have been reported regarding ligand-independent TNFR1 activation^50,62–68^. Depletion of ESCRT (endosomal sorting complex required for transport) proteins results in a clustering of internalized TNFR1 and LTβR at endosomes, initiating the NF-κB signaling cascade in a ligand-independent manner^50^. Importantly, EBOV VP40 interacts with multiple ESCRT proteins via its late domains or an uncharacterized mechanism and recruits ESCRT proteins from endosomes to the plasma membrane for efficient budding^69–71^. Thus, it is possible that the interaction between the host ESCRT machinery and VP40 affects the dynamics of TNFRSF at endosomes, thereby triggering the ligand-independent NF-κB signaling activation. The mapping analyses in our study strongly suggest that the domain spanning 21-44 aa within EBOV VP40 is a key candidate involved in this mechanism. Investigating this hypothesis by assessing the cellular distribution of TNFRSF in the presence of VP40, as well as conducting a comprehensive interactome analysis targeting the VP40 HVR, will be a focus of future research.

In summary, we propose a novel molecular mechanism of sustained pro-inflammatory activation mediated by EBOV VP40 and a previously unrecognized function for VP40 as a virulence determinant among ebolaviruses. Although previous reports have suggested that EBOV VP40 may regulate host cellular responses, including inflammatory and antiviral responses^72–74^, our study may offer a potential molecular mechanism for the induction of uncontrolled inflammatory responses linked to ebolavirus pathogenesis. Further investigation into the molecular basis of the host-EBOV interaction involved in EBOV-induced inflammatory activation may advance the development of therapeutic approaches targeting the interaction interface between host and viral proteins that trigger pathogenic, sustained inflammatory responses in severe EVD.

## Methods

### Cells and transfections

293 (CRL-1573), 293-TLR4/MD2 (BEI Resources), Huh7 (a kind gift from Dr. Yoshiharu Matsuura, Osaka University), HepG2 (ATCC HB-8065), and Vero E6 cells (ATCC CRL-1586) were maintained in DMEM supplemented with 10% FBS and 1% penicillin-streptomycin (PS) (growth medium) in 5% CO_2_ incubator at 37°C. Transient transfection was performed with Transit-LT1 (Mirus) according to the manufacturer’s instructions unless mentioned otherwise.

### Viruses and biosafety statement

EBOV (variant Mayinga), BDBV, and RESTV (variant Pennsylvania) were propagated in Vero E6 cells. Virus infectivity titers (focus-forming units, FFU) were determined by counting the number of infected cell foci using an indirect immunofluorescent antibody assay using a rabbit polyclonal anti-VP40 antibody as a primary antibody^75^, as previously described^76,77^. All work with infectious ebolaviruses was performed under biosafety level 4 conditions at the Rocky Mountain Laboratories Integrated Research Facility (Hamilton, Montana). Standard operating protocols were approved by the Institutional Biosafety Committee.

### Plasmids

EBOV (variant Mayinga) protein expression plasmids pCAGGs-NP, -VP35, -VP40, -GP, -VP30, -VP24 and -L were previously generated^75,78^ utilizing a pCAGGs vector possessing the cytomegalovirus enhancer fused to the chicken beta-actin promoter^79^. Additionally, VP40 derived from various EBOV variants (variant Kikwit and Makona-C07), as well as N-terminally FLAG-tagged VP40 from EBOV (variant Mayinga), SUDV (variant Gulu), BDBV, TAFV, and RESTV (variant Pennsylvania), and TRAF6 were cloned into a pCAGGs vector using standard cloning techniques.

A pNFκB-luc (AF053315.1) encoding a firefly luciferase gene driven by five copies of NF-κB response elements positioned upstream of a minimal promoter, and a pRL-TK (Promega) encoding a *Renilla* luciferase gene under the control of HSV-thymidine kinase promoter were utilized for dual-luciferase reporter (DLR) assays.

The following open reading frames were cloned into an expression vector under the control of the phosphoglycerate kinase promoter: IKK-2 and its mutant IKK-2 K44M (Addgene plasmids #11103 and #11104, gifts from Anjana Rao^80^); 3xHA-Iκβα and its mutants 3xHA-Iκβα-SS32/36AA (Addgene plasmids #21985 and #24143, gifts from Warner Greene^81^); TNFR1-YFP (Addgene plasmid #111209, gift from Johannes A. Schmid).

Single guide RNA (gRNA) (**Supplementary Table 1**) were cloned into a lentiCRISPR v2 (Addgene plasmid #52961, a gift from Feng Zhang^82^).

### Luciferase reporter assays

For NF-κB-responsive luciferase reporter assays with individual protein expression, 293 cells (6 x 10^4^ cells), Huh7 (5 x 10^4^ cells), and HepG2 (4 x 10^5^ cells) were seeded in 24-well plate 1 day before transfection. Cells were transfected with a pNFκB-luc (0.25 μg) and a pRL-TK (0.04 μg) together with 0.5 μg of expression plasmids unless mentioned otherwise. Cells were lysed using Passive Lysis Buffer (PLB) (Promega) at the indicated time points, and luciferase activities were measured using DLR System (Promega). The results are presented as relative light units (RLU), calculated as the ratio of firefly luciferase activity to *Renilla* luciferase activity.

For NF-κB luciferase reporter assays with virus infection, 293 cells (1 x 10^5^ cells) were seeded in 24-well plate 1 day before transfection. Cells were transfected with a pNFκB-luc (0.25 μg) and a pRL-TK (0.04 μg). Next day, cells were infected with ebolaviruses at a multiplicity of infection (MOI) of 1. After 1 h adsorption with tilting every 15 min, cells were washed once and 1 ml of DMEM supplemented with 3% FBS was added to the cells. Cells were lysed using PLB at 24, 48, and 72 hpi, and luciferase activities were measured using DLR System. Cell supernatants were harvested at 0, 24, 48, and 72 hpi, and used for virus titration.

### Western blotting

Cell lysates were prepared in either RIPA lysis buffer containing 1% NP-40, NE-PER Nuclear and Cytoplasmic Extraction reagents (Thermo Scientific), or PLB. Protease inhibitor cocktail (MilliporeSigma) and phosphatase inhibitor cocktail (Thermo Scientific) were added into lysis buffer accordingly. The total protein amount was determined using Pierce BCA Protein Assay Kits (Thermo Scientific). An equivalent amount of proteins was subjected to SDS-PAGE and proteins were transferred onto PVDF membranes. The primary antibodies are shown in **Supplementary Table 2**. Donkey anti-mouse and donkey anti-rabbit secondary antibodies conjugated to horseradish peroxidase (Jackson ImmunoResearch Laboratories, 1:25000) were used along with enhanced chemiluminescence substrate (Thermo Scientific).

### Nuclear and Cytoplasmic fractionation assay

293 cells (3.5 x 10^5^ cells) were seeded in 6-well plate 1 day before transfection. Cells were transfected with the indicated expression plasmids (2 μg) and were harvested at 24 hpt. NE-PER Nuclear and Cytoplasmic Extraction reagents was used according to the manufacturer’s instruction. The extracted fractions were utilized for protein detection by western blotting, with LaminA/C and β-tubulin serving as controls for nuclear and cytoplasmic protein, respectively.

### Gene silencing by siRNAs

siRNA-mediated gene silencing was performed using Dharmacon ON-TARGETplus SMARTpool siRNA. 293 cells (1 x 10^5^ cells) were seeded in 12-well plate. Immediately after cell seeding, cells were transfected using DharmaFECT 1 (Dharmacon) with a 25 nM siRNA targeting NF-κB p65 (RelA) subunit, GAPDH, or non-targeting negative control. Next day, the cells were transfected with a pNFκB-luc (0.5 μg) and a pRL-TK (0.08 μg) together with a pCAGGs-EBOV VP40 (1 μg). Forty-eight hours later, cells were harvested in PLB and cell lysates were used for DLR assay and western blotting.

### Gene knockout by CRISPR-Cas9

293 cells (3 x 10^5^ cells) were seeded in 6-well plate 1 day before transfection. Cells were transfected with a lentiCRISPR v2 encoding gRNA targeting Iκβα, TNFR1, or TNFα (0.3 μg). Cells transfected with lentiCRISPR v2-Iκβα or -TNFR1 were passaged several times under 1 μg/ml of puromycin selection. Monoclonal cell lines were established by limiting dilution method and genomic DNA was extracted from each cell line using QIamp DNA Mini Kit (Qiagen). The extracted DNA was then utilized as the template for PCR amplification, generating a fragment of approximately 300 bp that includes CRISPR editing site. PCR amplicon was employed for Sanger sequencing, and CRISPR editing were determined by using the Synthego ICE Analysis tool v3.0, western blotting, and flow cytometry (**Fig 6b**, **Supplementary Fig. 10, 11**).

### Flow cytometry

293 cells or TNFα knockout 293 cells (3.5 x 10^5^ cells) were seeded in 6-well plate 1 day before transfection, and were transfected with a pCAGGs-EBOV VP40 (2 μg). At 72 hpt, cells were treated with BD GolgiPlug Protein Transport Inhibitor (Brefeldin A) (BD Biosciences) at a final concentration of 1:1000 for 6 hours followed by fixation with 2% paraformaldehyde for 30 min. Cells were permeabilized with Intracellular Staining Permeabilization Wash Buffer (BioLegend) and then stained with 3 μl of PE-anti human TNFα antibody (BioLegend) in a 100μl of volume for 30 min. The samples were measured using BD LSR II Flow Cytometer. All flow cytometry data were analyzed using Flow Jo software.

### TNFα neutralization assay

293 cells (6 x 10^4^ cells) were seeded in 24-well plate 1 day before transfection. For TNFα treatment as a control, cells were transfected with a pNFκB-luc (0.25 μg) and a pRL-TK (0.04 μg). Next day, recombinant human TNFα (R&D Systems) (0.05 ng) and purified anti-human TNFα antibody (BioLegend) (12.5 μg) were mixed into 0.5 ml of growth medium and the complex was incubated for 1 hour at room temperature. The supernatant from plasmid-transfected cells was then removed and replaced with medium containing the TNFα -TNFα NAb complex. Cell supernatant and cell lysates were harvested at 6 hours post-treatment. For VP40 transfection, cells were transfected with a pNFκB-luc (0.25 μg), a pRL-TK (0.04 μg), and pCAGGs-EBOV VP40 (0.5 μg). Immediately after transfection, TNFα NAb (0.05 ng) was added into the medium of transfected cells. Cell supernatant and cell lysates were harvested at 48 hours post-treatment. Cell supernatant was stored in -80°C until use.

For transferring supernatants, 293 cells (6 x 10^4^ cells) were seeded in 24-well plate 1 day before transfection and were transfected with a pNFκB-luc (0.25 μg) and a pRL-TK (0.04 μg). Next day, the supernatant was removed and replaced with previously harvested cell supernatant. Cell lysates were harvested at 8 hours post-treatment and were used for DLR assays.

### qRT-PCR

293 cells (4 x 10^4^ cells) were seeded in 6-well plate 1 day before transfection. Cells were transfected with a pCAGGs-EBOV VP40 (2 μg) and were lysed using Trizol at 24 hpt for RNA extraction. RNA was purified using Direct-zol RNA MiniPrep Kit (Zymo Research) with in-column DNAseI treatment according to manufacturer’s directions. The RNA of IL-8 (CXCL8), TNF-α, and MIP-1β (CCL4) was quantified by qRT-PCR using iTaq Universal Probes One-Step Kit (BioRad). Each reaction used 0.1 μg of RNA, 0.2 μM of forward and reverse primers, and 0.1 μM or 0.05 μM of probe with 5’ 6FAM fluorophore and 3’ BlackBerry quencher (BBQ) (TIB Molbiol) for target cytokine RNA and control GAPDH RNA, respectively. The primer/probe sequences are shown in **Supplementary Table 3**. Cycling conditions were as follows: 10 min initial reverse transcription at 50°C, 2 min initial denaturation/activation at 95°C, followed by 40 cycles of 5 sec denaturation at 95°C and 10 sec annealing/extension step. Delta-delta *C*_t_ values were used to determine their relative expression as fold changes. qPCR was performed using the CFX384 Touch Real-Time PCR Detection System (BioRad), and data were analyzed with BioRad CFX Manager 3.1.

### RNA-seq and transcriptomic analysis

293 cells (2 x 10^5^ cells) were seeded in 12-well plate 1 day before infection. Cells were infected with EBOV (variant Mayinga), RESTV (variant Pennsylvania) at MOI of 1, or mock-infected with culture medium. The infection experiment was independently repeated three times (n = 3). Cells were inactivated using Trizol at 24 and 48 hpi, and samples were sent to the University of Saskatchewan where the RNA extraction was performed with the Zymo Direct-Zol kit per the manufacturer’s protocol. RNA quality control was performed using an Agilent Bioanalyzer 2100 and samples were sent to the Saskatchewan Global Institute for Food Security (GIFS), where library prep was performed using the Illumina TruSeq Stranded mRNA Library kit and libraries were run on an Illumina NovaSeq6000.

Alignments and differential expression analysis were performed using an established pipeline. The sequence read quality of all 36 samples was examined using Fastqc v0.11.9^83^. The paired-end fastq reads were mapped to the human reference genome assembly GRCh38^84^ (ensemble release-109) using STAR v2.7.10b^85^. The genome index was generated using parameter “--sjdbOverhang 150” in STAR using reference genome and its annotation in GTF format. The raw counts were generated from the alignment files using the Rsubread v2.12.3 Bioconductor package^86^ in R v4.2.2. The resulting read counts were imported in R and genes were filtered based on expression level (to keep only rows that have a count of 15 or higher in at least 6 samples) and differential expression analysis was performed using Bioconductor package DESeq2 v1.38.3^87^. Differentially expressed genes (DEGs) were identified using Wald test in DESeq2. To control the false discovery rate (FDR), p values were adjusted by applying the Benjamini-Hochberg (BH) method at an FDR cut-off of 5% (i.e., alpha=0.05) during differential expression analysis. Multidimensional scaling (MDS) plot of differentially expressed genes (filtered using adjusted *P*-value <0.01 and log2 fold change >1.5 and -1.5) was generated in R using ggplot2, vsn, and tidyverse packages.

For functional analysis, DEGs were uploaded to IPA (QIAGEN Bioinformatics), a manually curated database of experimentally demonstrated intermolecular relationships from the peer-reviewed literature for human, rat, and mouse experimental systems. The IPA Knowledgebase is updated on a quarterly basis and these analyses were performed with the most recent update (spring 2024) at the time of writing. The thresholds (fold change relative to time-matched, mock-infected controls > |1.5|, adjusted *P*-value < 0.05) were applied prior to running IPA Core Analysis.

Gene set enrichment analysis (GSEA) was used to analyze DEGs using hallmark gene sets available from the most recent release of the Human Molecular Signatures Database (MSigDB)^88^ and clusterProfiler^89^ v4.10.1 in R v4.3.3. The thresholds (fold change relative to time-matched, mock-infected controls > |1.5| (log2FC > |0.585|), adjusted *P*-value < 0.05) were applied prior to GSEA as described above. Pathways that contained 10 or more DEGs were selected for analysis. Dot plots were generated using clusterProfiler version 4.10.1.

### Cytokine quantification

For protein expression, 293 cells (2 x 10^5^ cells) were seeded in 12-well plate 1 day before transfection. Cells were transfected with a pCAGGs-EBOV VP40 or -empty vector (1 μg) and supernatants were harvested at 24, 48, and 72 hpt.

For virus infection, 293 cells (2 x 10^5^ cells) were seeded in 12-well plate 1 day before infection. Cells were infected with EBOV, BDBV, or RESTV at MOI of 1 and supernatants were harvested at 24, 48, and 72 hpi. Cytokine levels were quantified using Bioplex human cytokine/chemokine 12-plex assay (BioRad) with Luminex Magpix instrument. Analyzed cytokines were as follows: TNF-α, IL-6, IL-1RA, IL-1β, IL-17 (IL-17A), IFN-γ, IL-8 (CXCL8), MIP-1β (CCL4), PDGF-BB, MCP-1 (CCL2), IP-10 (CXCL10), RANTES (CCL5).

### Statistical analysis

All experiments were performed as at least three independent biological replicates. Statistical analyses were performed using either the t-test or one-way ANOVA with GraphPad Prism 10 version 10.0.2.

## Data availability

All data generated from this study are available within the article and Supplementary Information files. Raw FASTQ files for RNA-seq have been deposited to the NCBI Sequence Read Archive and can be accessed via BioProject PRJNA1040271.

## Code availability

The step-by-step analysis, codes and parameters used for the transcriptomics data analysis are present at https://github.com/rasmussen-lab/Ebola-VP40.

## Acknowledgements

We thank Dr. Logan Banadyga from the University of Manitoba, and Dr. Andrea Marzi and Dr. Atsushi Okumura from the Rocky Mountain Laboratories, NIAID, NIH for their insightful discussions and for providing the plasmids used for NF-κB-responsive luciferase reporter assays. We are also grateful to Friederike Feldmann from the Rocky Mountain Laboratories, NIAID, NIH for her invaluable assistance with BSL-4 work. We also thank Carla M. Weisend from Mayo Clinic for her exceptional support in managing the project budget and overseeing laboratory operations throughout this research. We acknowledge Dr. Jie Sun and Dr. Chaofan Li from the University of Virginia for their crucial support in analyzing the preliminary transcriptomic data of VP40-expressing cells. This work was supported in part by the National Institute of Allergy and Infectious Diseases [R01 AI134937] and the Division of Intramural Research, NIAID, NIH. The opinions, interpretations, conclusions, and recommendations are those of the authors and are not necessarily endorsed by the NIH.

## Author contributions

Conceptualization: H.E., S.Y.; Methodology: S.Y., H.E., A.R., S.B., S.L.; Software: Z.M., R.S., S.S., A.R., V.S.; Validation: S.Y., L.W.; Formal analysis: Z.M., R.S., S.S., A.R., V.S.; Investigation: S.Y., L.W., K.M., S.L., B.Z.; Resources: H.E., M.B., A.R., S.B.; Data curation: Z.M., R.S., S.S., A.R., V.S.; Writing -Original Draft: S.Y., H.E., A.R.; Writing - Review & Editing: all authors; Visualization: S.Y., Z.M., R.M., V.S.; Supervision: H.E., A.R., M.B., S.B.; Project administration: S.Y.; Funding acquisition: H.E., M.B., A.R.

---

**Supplementary Fig. 1:**
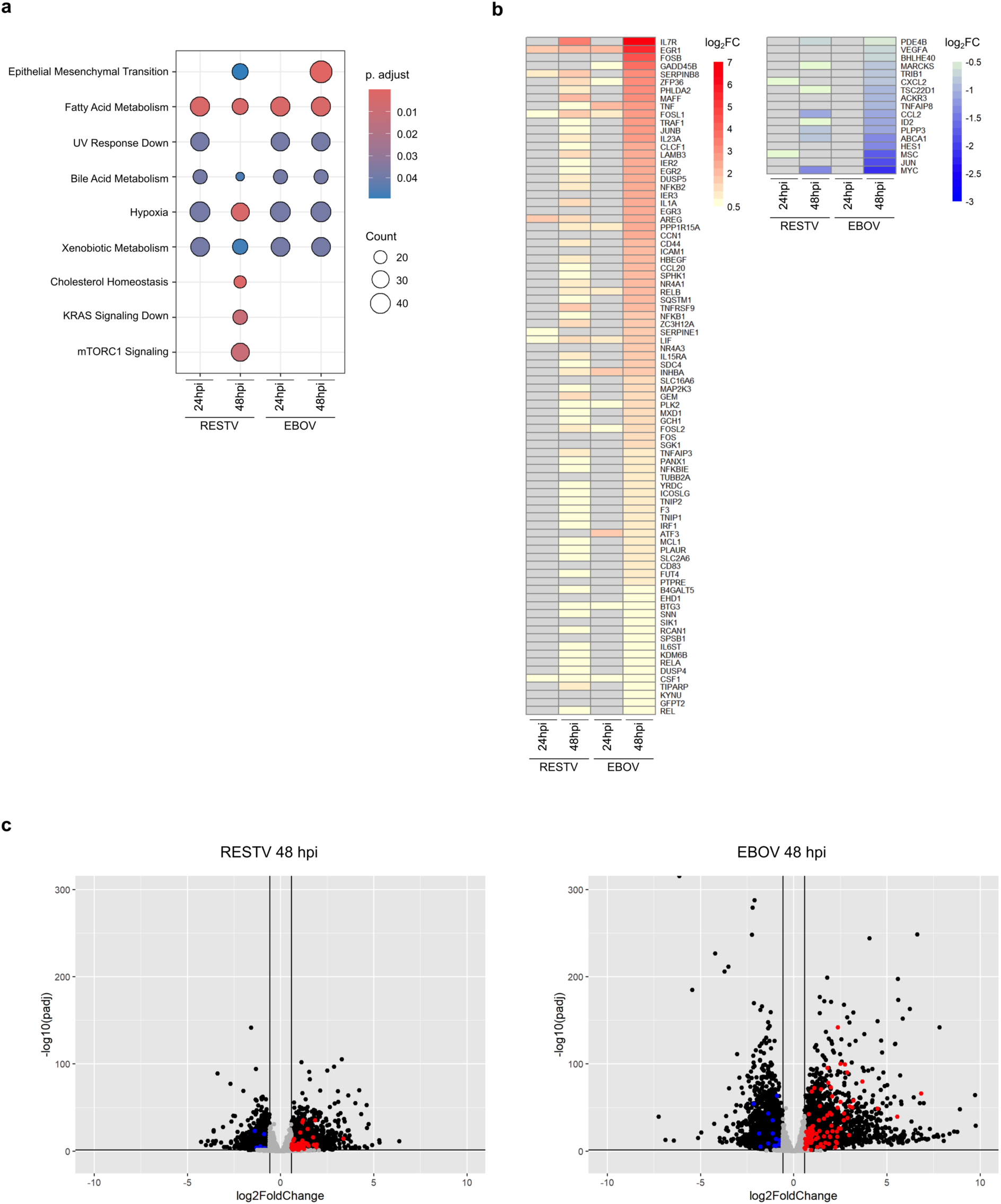
Transcriptomic analyses of Hallmark ‘TNFA Signaling via NFKB’ gene set using RNA-seq results from EBOV, or RESTV-infected 293 cells. **a** Dot plot showing GSEA hallmark pathways that contained 10 or more downregulated DEGs. Gene sets were sorted to only include genes that reached criteria (fold change relative to time-matched, mock-infected controls < 1.5, adjusted *P*-value < 0.05) within the EBOV 48 hpi dataset. Further details on GSEA are provided in the Materials and Methods section. **b** Heatmaps for upregulated DEGs (left) and downregulated DEGs (right) generated using the Hallmark ‘TNFA Signaling via NFKB’ (M5890)^90^ gene set available from MSigDB. Gene sets were sorted to only include genes that reached criteria (fold change relative to time-matched, mock infected controls > |1.5| [log_2_FC > |0.585|], adjusted *P*-value < 0.05) within the EBOV 48 hpi dataset. Gray color indicates genes that did not reach DE criteria in given dataset. Heatmaps were generated using the R package pheatmap v1.0.12^91^. **c** Volcano plots showing DEGs in RESTV and EBOV 48 hpi datasets. Red dots indicate upregulated and blue dots indicate downregulated DEGs within the ‘TNFα Signaling via NF-κB’ GSEA Hallmark pathway. Gray dots indicate genes that did not meet the DE criteria. Black dots indicate all other DEGs not within the ‘TNFα Signaling via NF-κB’ GSEA Hallmark pathway. Vertical lines indicate log_2_FC cutoffs, horizontal line indicates adjusted *P*-value cutoff. Plots were generated using the R package ggplot2^92^ v3.5.1.

**Supplementary Fig. 2:**
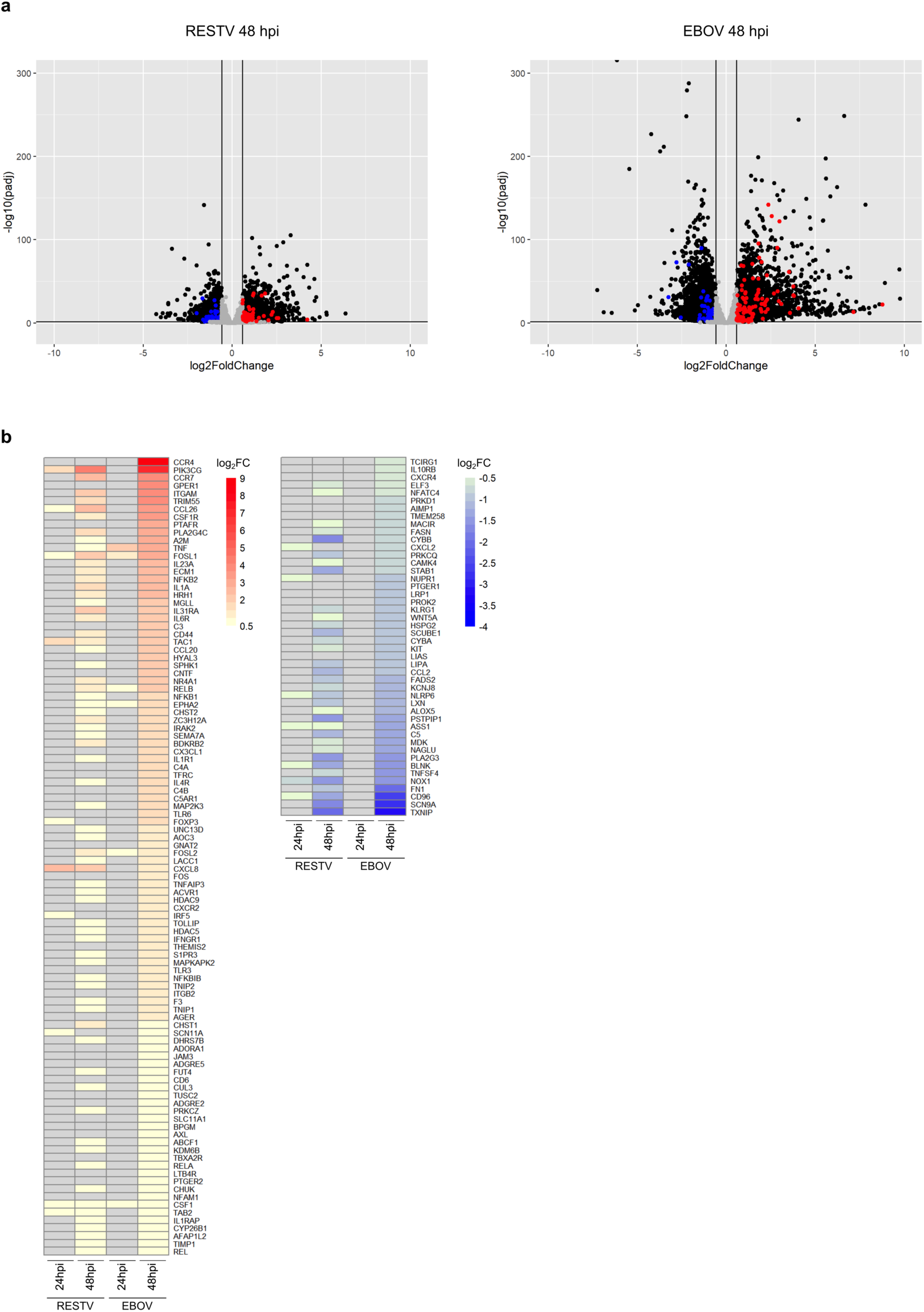
Transcriptomic analyses of ‘Inflammatory Response (GO:0006954)’ gene set from AmiGO 2 using RNA-seq results from EBOV-or RESTV-infected 293 cells. **a** Volcano plots. Red dots indicate upregulated and blue dots indicate downregulated DEGs within the ‘Inflammatory Response (GO:0006954)’ from AmiGO 2 sorted to include only protein coding transcripts from *Homo sapiens*. Gray dots indicate genes that did not meet the DE criteria. Black dots indicate all other DEGs not within the ‘Inflammatory Response (GO:0006954)’ gene set. Vertical lines indicate log_2_FC cutoffs, horizontal line indicates adjusted *P*-value cutoff. Plots were generated using the R package ggplot2^92^ v3.5.1. **b** Heatmaps for upregulated DE genes (left) and downregulated DE genes (right) generated using the ‘Inflammatory Response (GO:0006954)’ gene set. Gene sets sorted to only include genes that reached criteria (absolute fold change relative to time-matched, mock infected controls > 1.5 [log2FC > 0.585], adjusted *P*-value < 0.05) within the EBOV 48 hpi dataset. Grey color indicates genes that did not reach DE criteria in given dataset. Heatmaps were generated using the R package pheatmap v1.0.12^91^.

**Supplementary Fig. 3:**
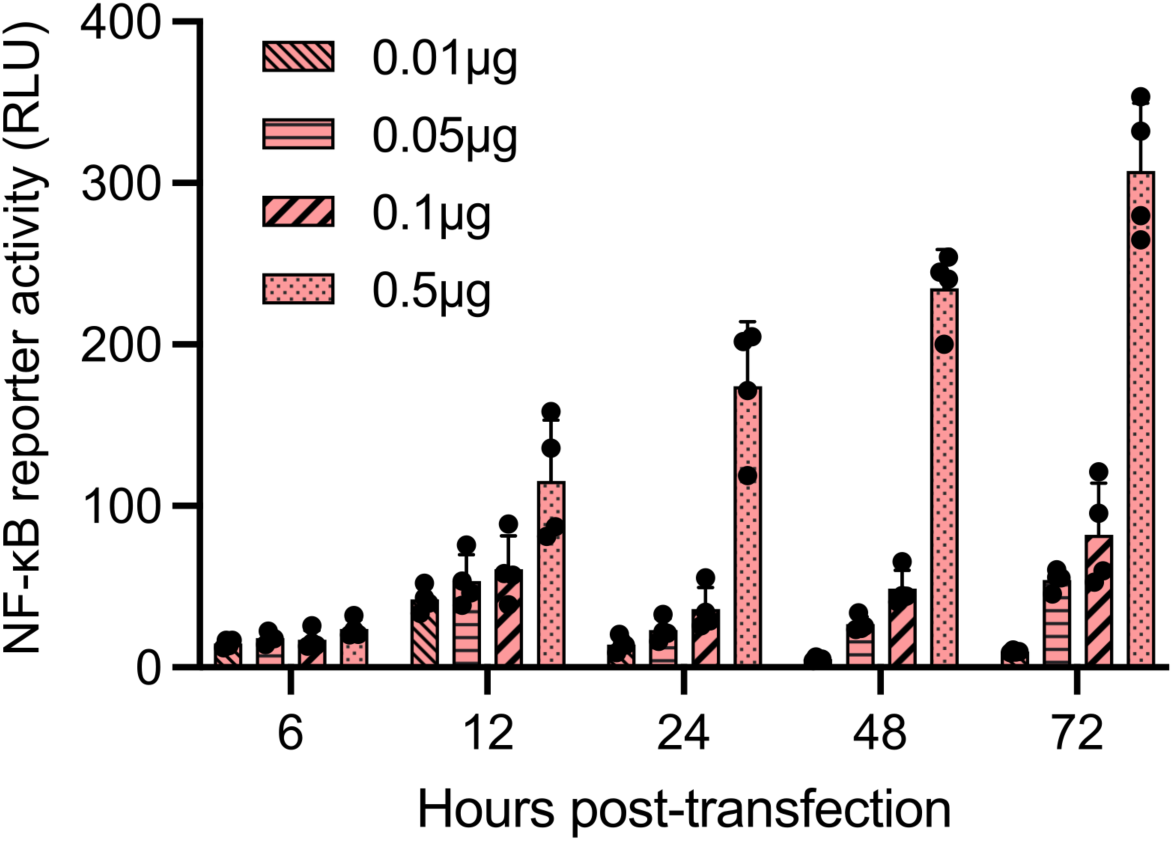
Time-and dose-kinetics of NF-κB-responsive reporter activity in 293 cells expressing EBOV VP40. Data are shown with mean ± SD (n = 3 independent biological replicates).

**Supplementary Fig. 4:**
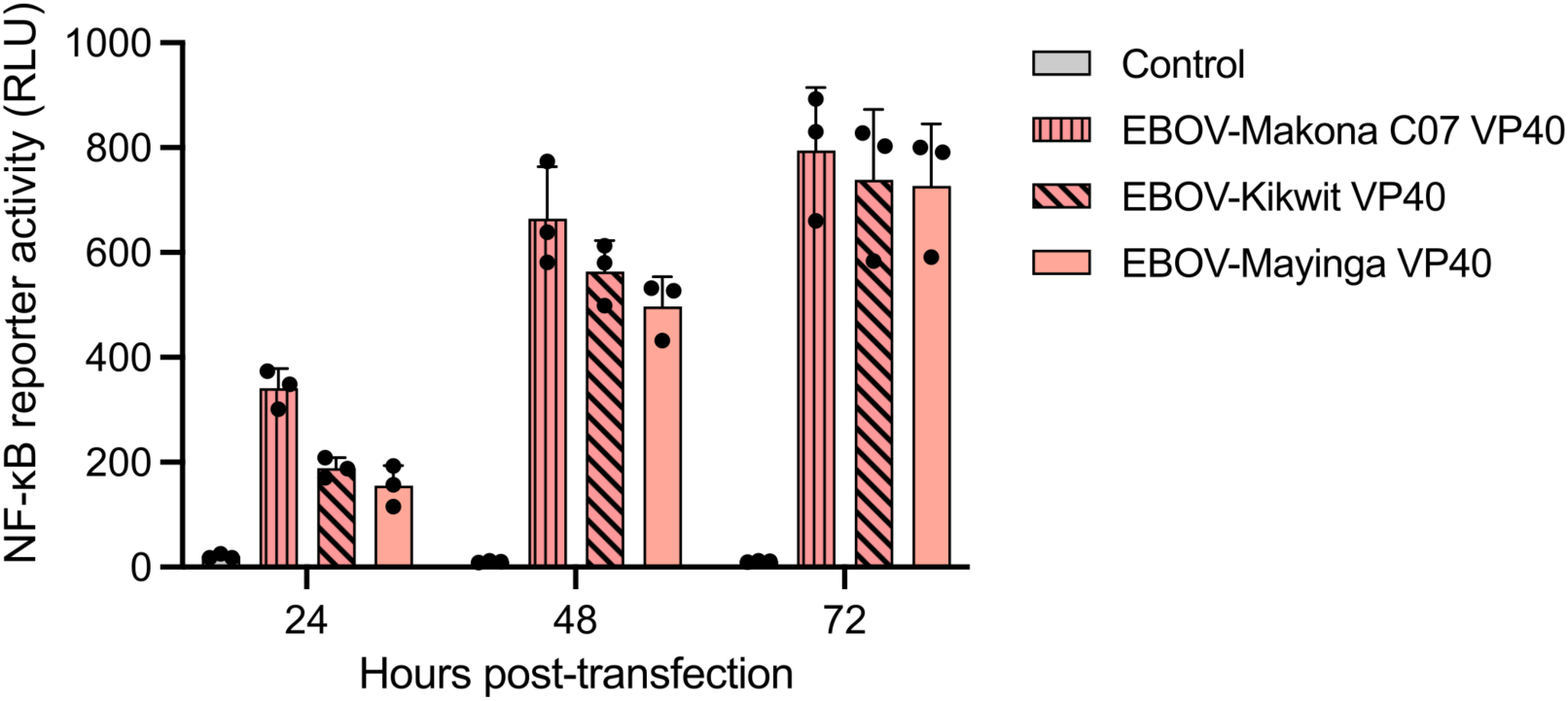
NF-κB-responsive luciferase reporter activity in 293 cells expressing VP40 derived from EBOV variant Mayinga, Kikwit, or Makona-07. Data are shown with mean ± SD (n = 3 independent biological replicates).

**Supplementary Fig. 5:**
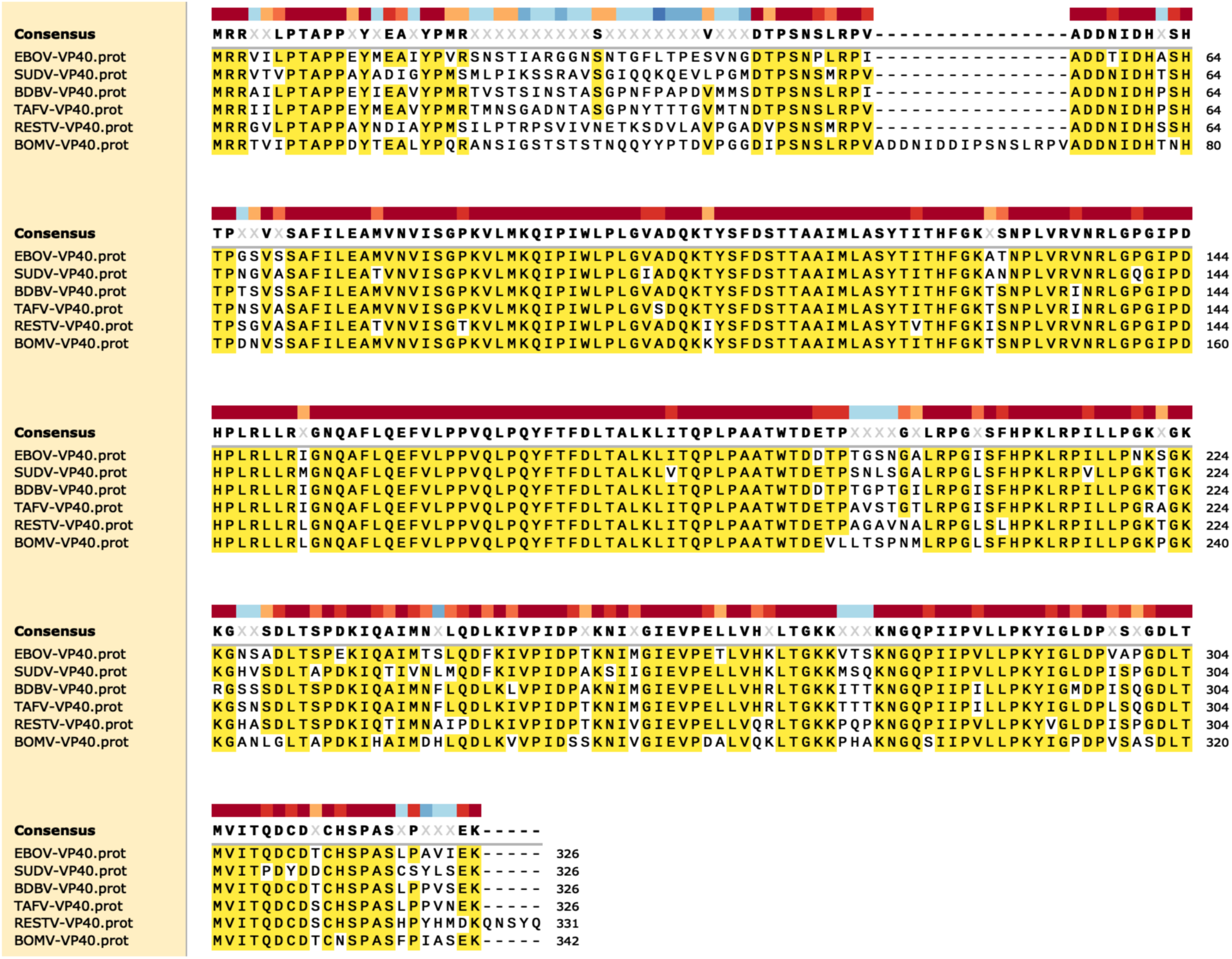
Amino acid sequence alignment of VP40 from six different ebolaviruses. Amino acid sequences were aligned using SnapGene version 7.2.1. Fully conserved amino acids across VP40 from six ebolaviruses are highlighted in yellow, with sequence conservation indicated by colored blocks.

**Supplementary Fig. 6:**
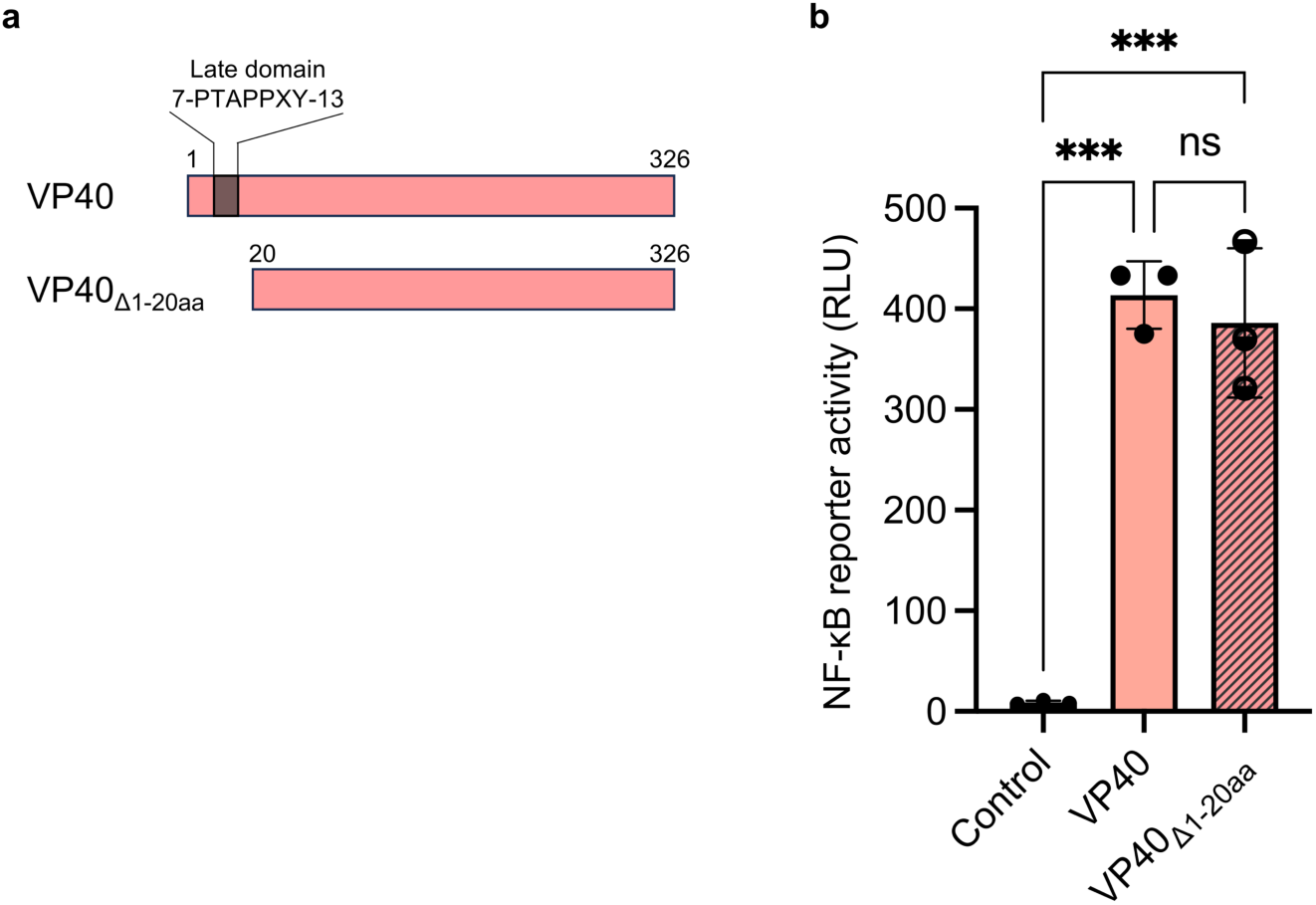
Examination of the importance of two classical late domains in VP40-mediated NF-κB activation. **a** Diagram of EBOV VP40 and a mutant with the first 20 amino acids deleted (VP40_Δ1-20aa_). **b** NF-κB-responsive luciferase reporter activity in 293 cells expressing either EBOV VP40 or VP40_Δ1-20aa_ at 72 hpt. Data are shown with mean ± SD (n = 3 independent biological replicates). ns > 0.05, \*\*\**P* ≤ 0.001; ordinary one-way ANOVA.

**Supplementary Fig. 7:**
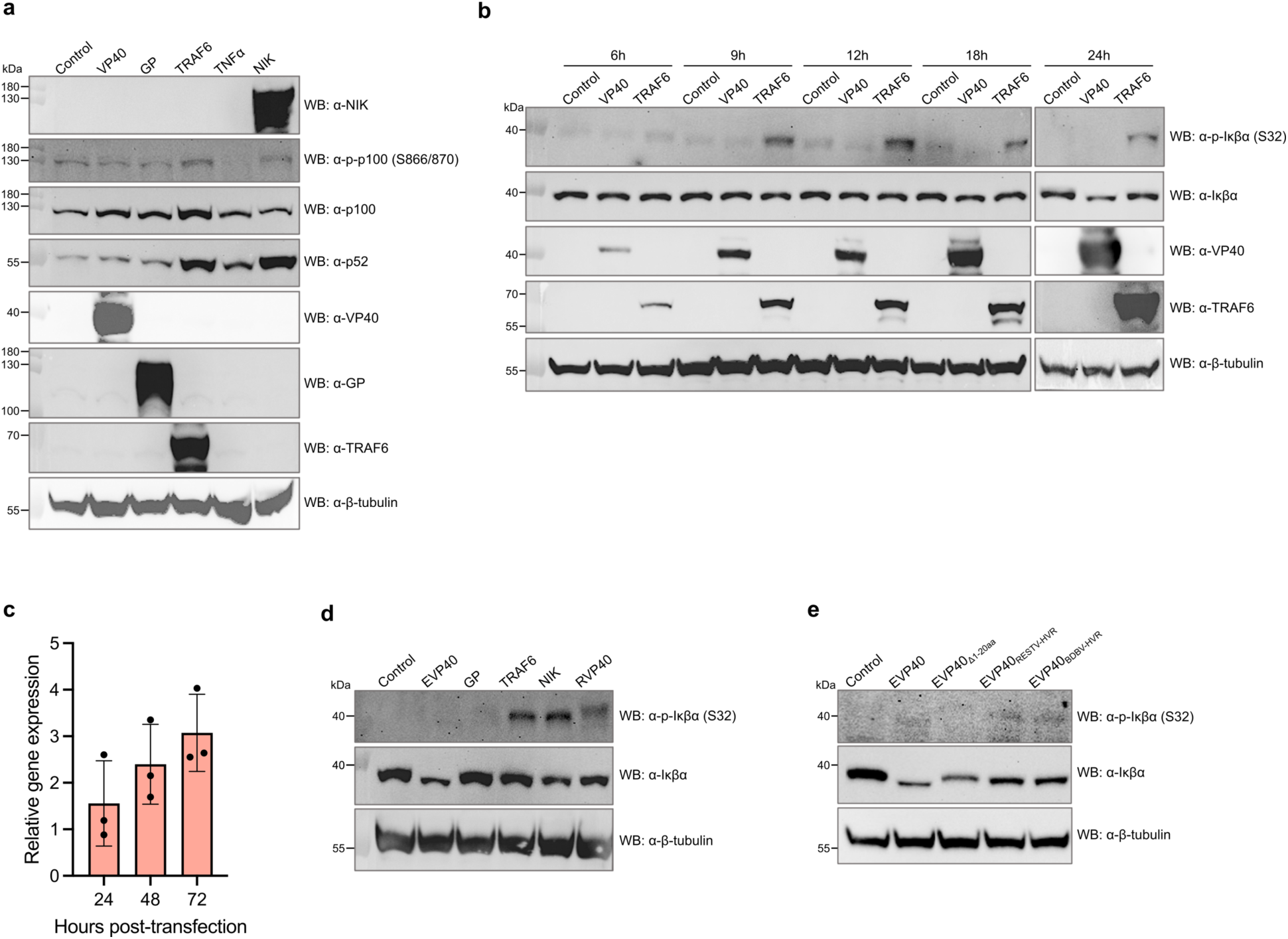
Examination of activation status of host proteins in NF-κB signaling pathways in the presence of EBOV VP40. **a** Western blotting for molecules involved in the non-canonical NF-κB pathway. 293 cells were transfected with plasmid EBOV VP40, EBOV GP, TRAF6, or NIK and harvested at 24 hpt or treated with 15 ng/ml of TNFα for 1 hour and then harvested. **b** Western blotting for phosphorylated-Iκβα or Iκβα in 293 cells expressing EBOV VP40 or TRAF6. **c** Quantification of Iκβα mRNA in EBOV VP40-expressing 293 cells. Extracted RNA was reverse transcribed using OligodT primer, and cDNA was used for qRT-PCR with iTaq Universal Probes Supermix (BioRad), Hs00355671_g1 NFKBIA (Thermo Scientific) and Hs00355671_g1 NFKBIA (Thermo Scientific). GAPDH was used as a reference gene (the primer/TaqMan probe sequences are shown in **Supplementary Table 3)**. Delta-delta *C*_t_ values were used to determine their relative expression as fold changes. Details on RNA preparation are provided in the Materials and Methods section. **d** Western blotting for phosphorylated-Iκβα or Iκβα in 293 cells expressing EBOV VP40, EBOV GP, TRAF6, NIK or RESTV VP40 at 24 hpt. **e** Western blotting for phosphorylated-Iκβα or Iκβα in 293 cells expressing EBOV VP40, EBOV VP40_Δ1-20aa_, or chimeric VP40 proteins where the HVR of EBOV VP40 was replaced with that of RESTV or BDBV (EVP40_HVR-RESTV_ and EVP40_HVR-BDBV_) at 24 hpt. Control: Empty vector transfected. For **c**, data are shown with mean ± SD (n = 3 independent biological replicates).

**Supplementary Fig. 8:**
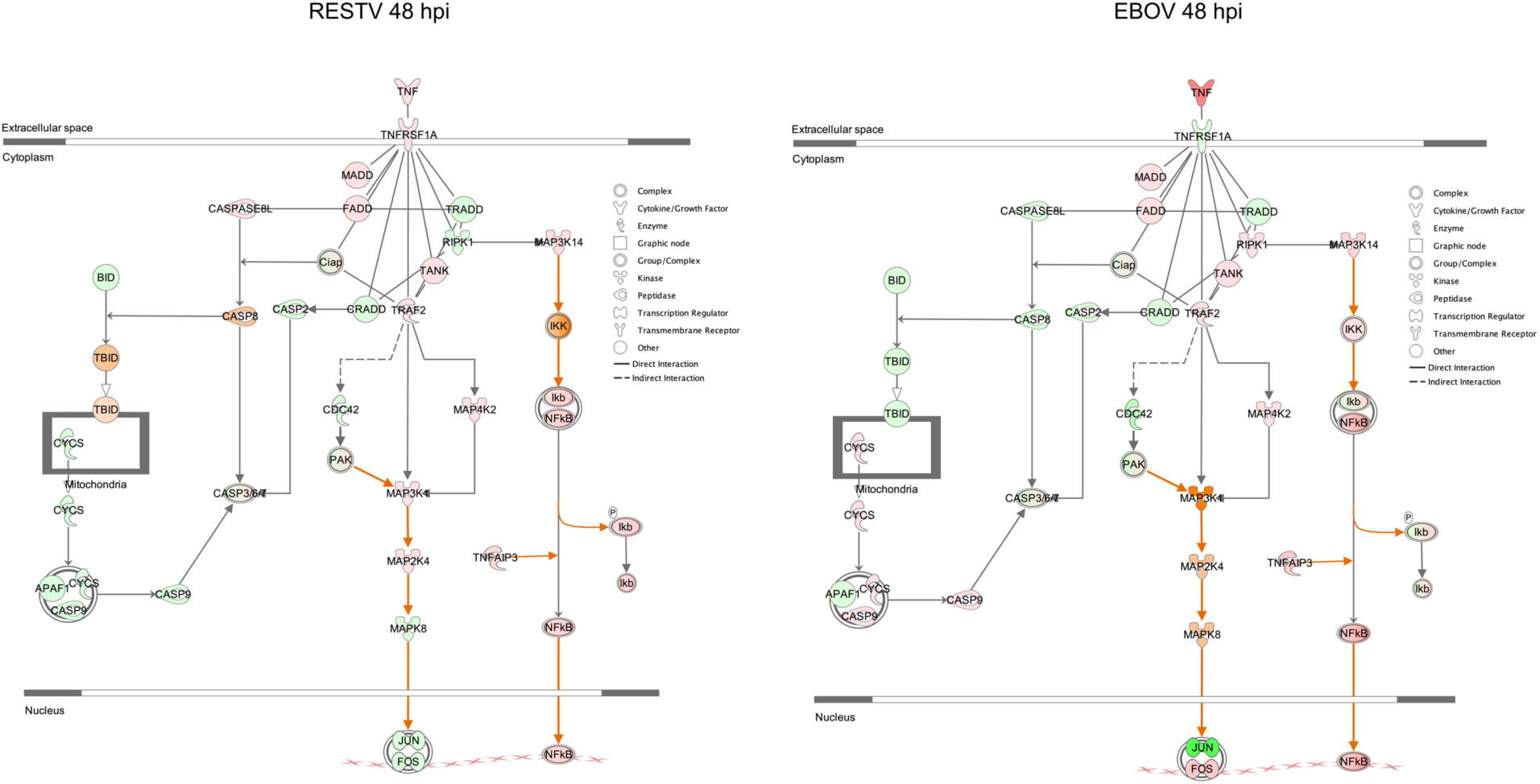
IPA TNFR1 Signaling Canonical Pathway overlaid with RNA-seq gene expression data from EBOV-or RESTV-infected 293 cells. Red and green shadings indicate positive fold change (upregulation) and negative fold change (downregulation) relative to the time-matched mock-infected controls, respectively. Orange shading indicates predicted activation. Darker red or green shading indicates that genes met DE criteria (fold change > |1.5|, adjusted *P*-value < 0.05), while lighter shading indicates that they did not. Darker orange shading indicates a higher z-score. Orange and blue lines indicate predicted activation and inhibition, respectively. Gray shading indicates insufficient information to predict molecular activity. Solid lines indicate known direct interactions. Arrowheads or line caps indicate directionality.

**Supplementary Fig. 9:**
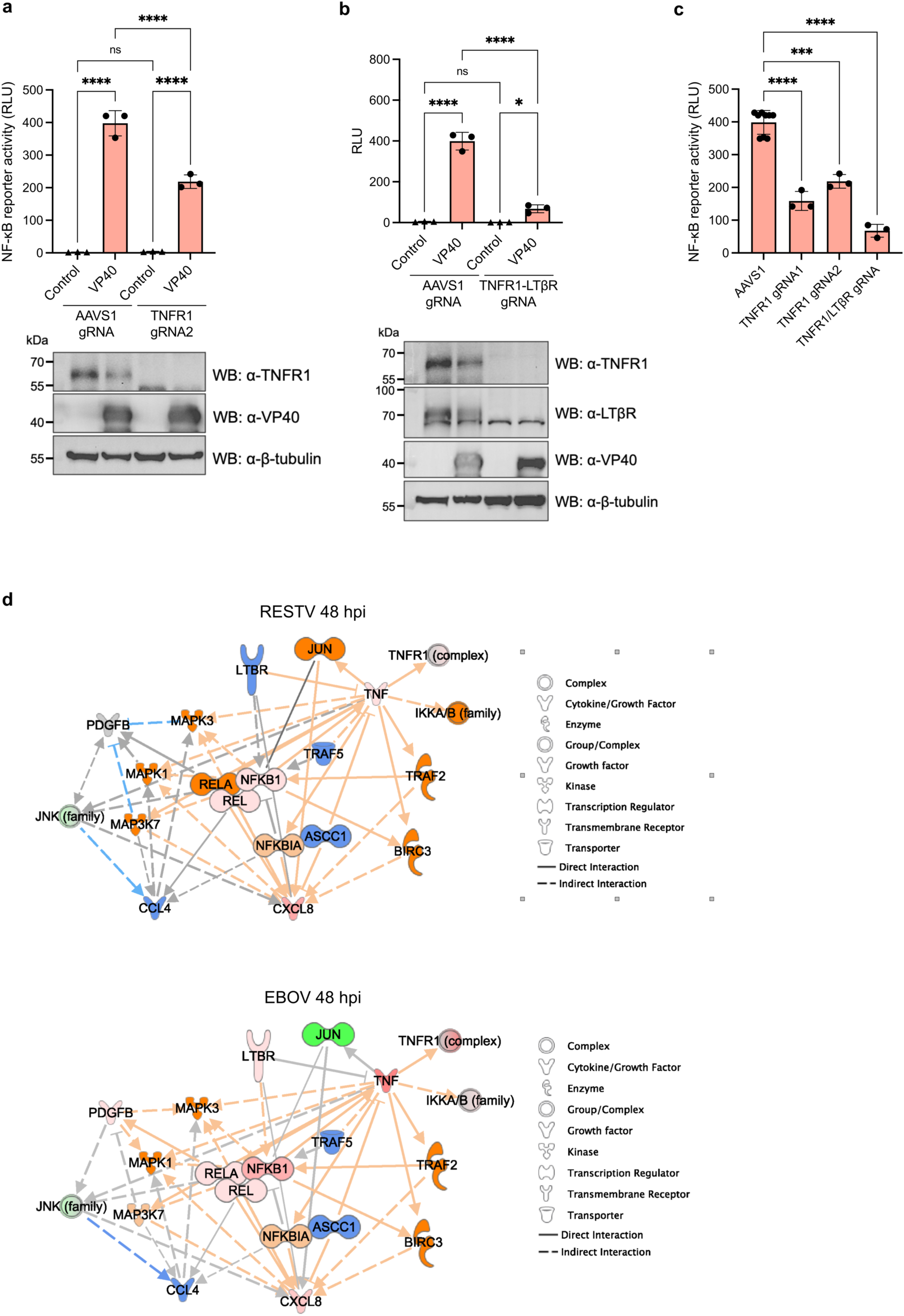
Assessment of contribution of TNFR1 to NF-κB activation induced by EBOV VP40. **a** NF-κB-responsive reporter activity in TNFR1-knockout (gRNA 2) 293 cells expressing EBOV VP40 at 48 hpt. **b** NF-κB-responsive luciferase reporter activity in TNFR1/LTβR-knockout 293 cells expressing EBOV VP40 at 48 hpt. **c** Summary of NF-κB-responsive reporter activity in TNFR1 or TNFR1/LTβR-knockout 293 cells expressing EBOV VP40 at 48 hpt. **d** IPA custom network generated with RNA-seq result from EBOV-or RESTV-infected 293 cells. Molecules associated with EBOV VP40-mediated inflammation-associated genes were selected to generate networks. NF-κB subunits (REL, RELA, NFKBI) are clustered at the network center as they directly interact and form a complex. Red and green shadings indicate upregulated and downregulated genes meeting DE criteria, respectively (fold change > |1.5|, adjusted *P*-value < 0.05). Orange and blue shadings indicate predicted activation and inhibition, respectively. Gray shading indicates insufficient information to predict molecular activity. Solid and dashed lines indicate known direct interactions and indirect interactions, respectively. Arrowheads or line caps indicate directionality. Control: Empty vector transfected. For **a**-**c**, data are shown with mean ± SD (n = 3 independent biological replicates). ns > 0.05, \**P* ≤ 0.05, \*\*\**P* ≤ 0.001, \*\*\*\**P* ≤ 0.0001; ordinary one-way ANOVA.

**Supplementary Fig. 10:**
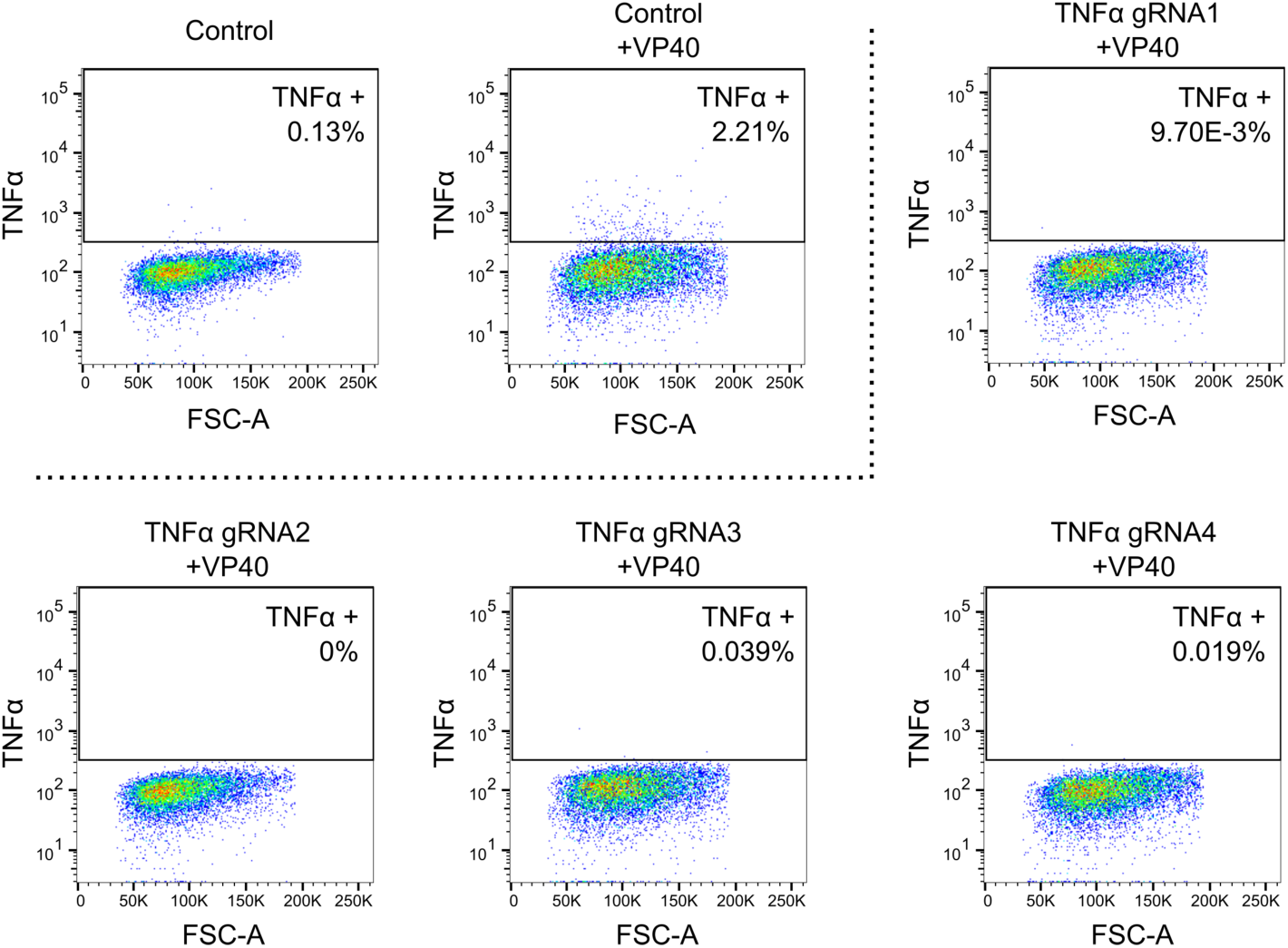
Quantification of intracellular TNFα in TNFα-knockout 293 cells expressing EBOV VP40 by flow cytometry. Control: CRISPR-Cas9 targeting AAVS1.

**Supplementary Fig. 11:**
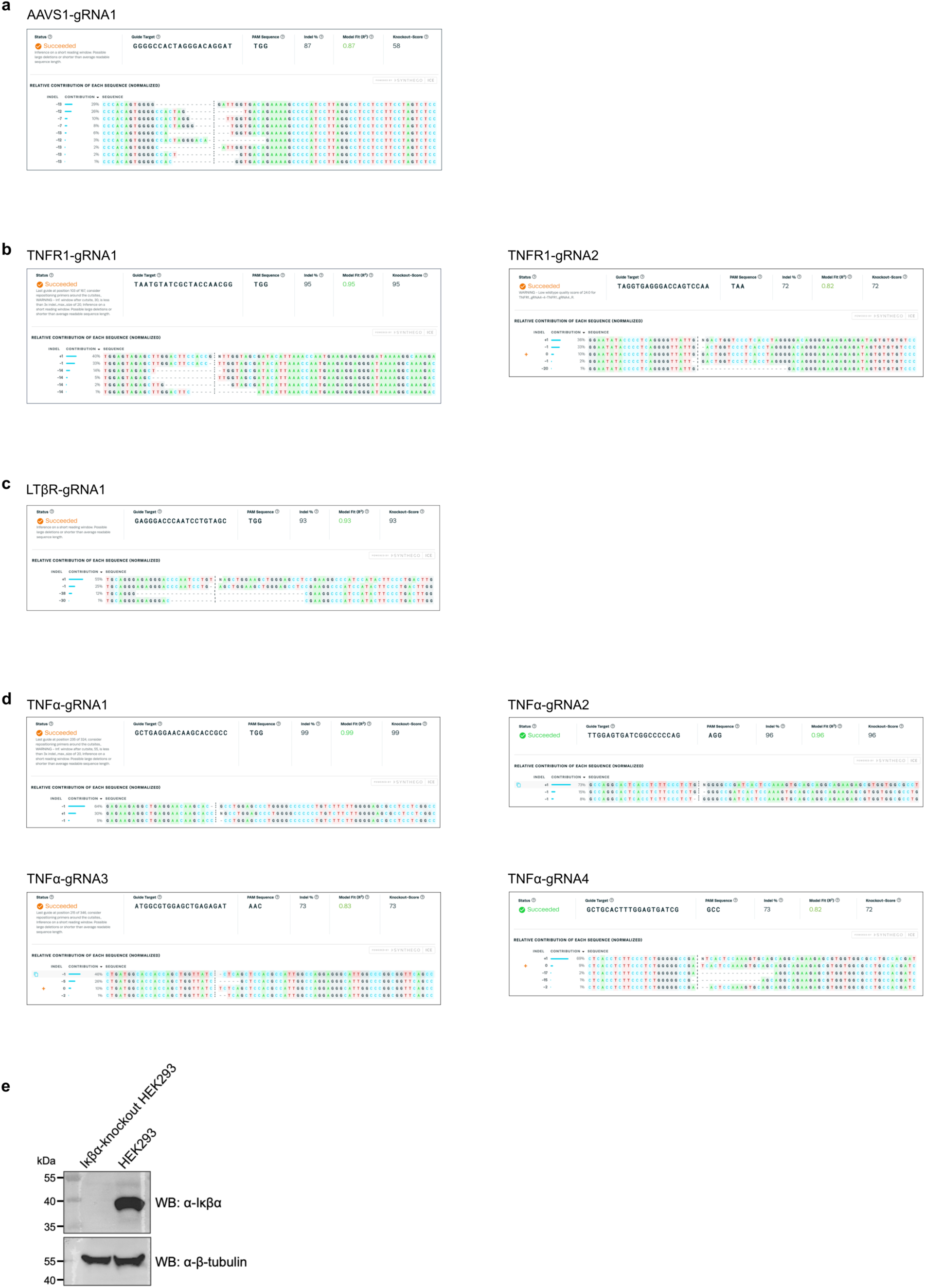
Confirmation of CRISPR editing. **a** Synthego ICE analysis for CRISPR editing targeting **a** AAVS1, **b** TNFR1, **c** LTβR, **d** TNFα. **e** Western blotting for Iκβα in Iκβα-knockout 293 and wild-type 293 cells.

**Supplementary Table 1.**
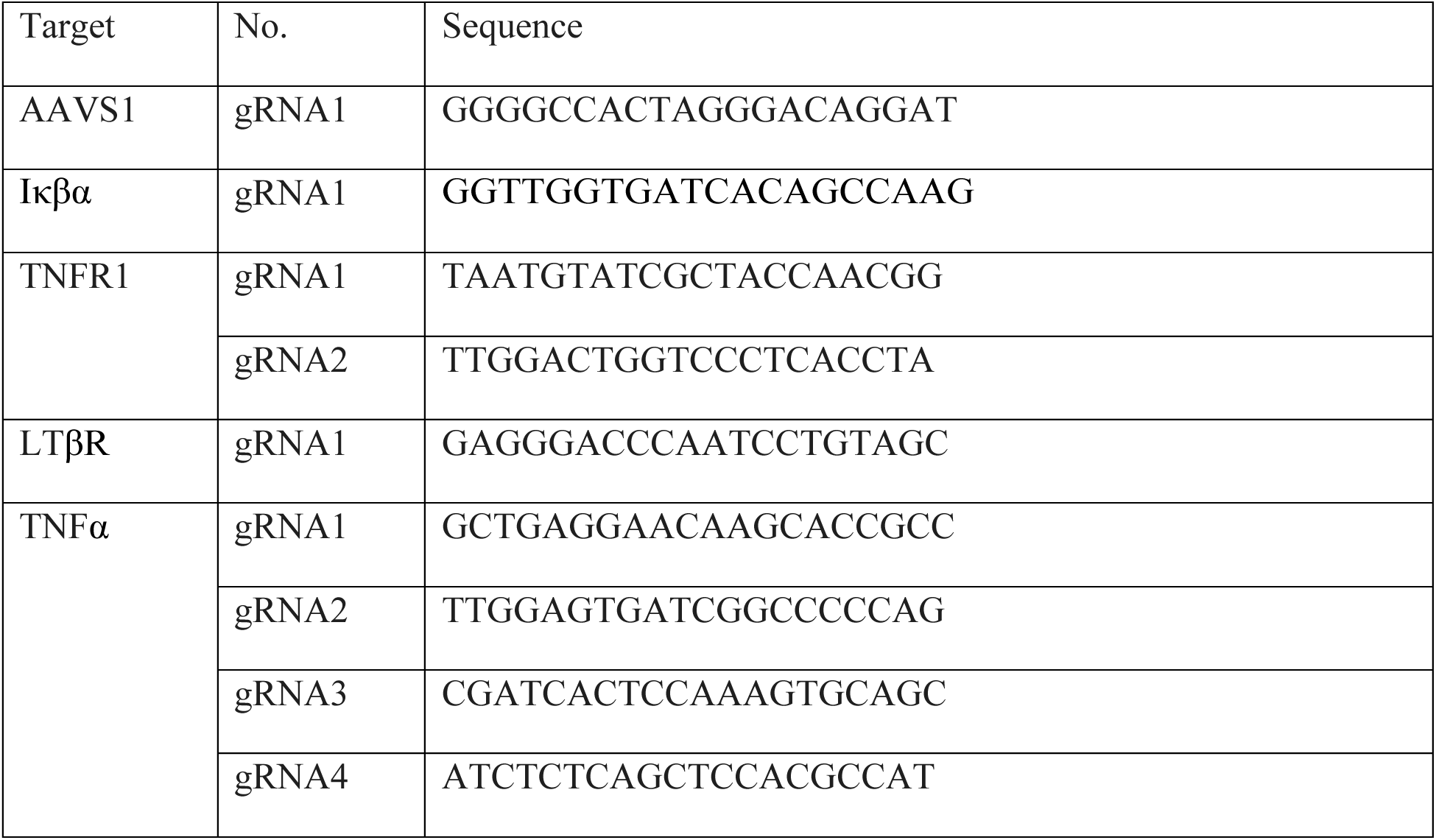
Sequences of gRNA.

**Supplementary Table 2.** List of primary antibodies used for western blotting in this study.

| Target protein | Company | Catalog number | Dilution |
| --- | --- | --- | --- |
| EBOV VP40 | IBT BioServices | 0201-017 | 1:2000 |
| EBOV GP | IBT BioServices | 0301-015 | 1:1000 |
| FLAG | MilliporeSigma | F7425 | 1:2000 |
| β-tubulin | Abcam | ab6046 | 1:2000 |
| TRAF6 | Cell Signaling Technology | 8028 | 1:1000 |
| LaminA/C | Cell Signaling Technology | 4777 | 1:1000 |
| GAPDH | Santa Cruz Biotechnology | sc-47724 | 1:500 |
| TLR4 | Cell Signaling Technology | Abcam | 1:1000 |
| p65 | Cell Signaling Technology | 4764 | 1:1000 |
| RelB | Cell Signaling Technology | 10544 | 1:1000 |
| c-Rel | Cell Signaling Technology | 12707 | 1:1000 |
| NF-κB1 p105/p50 | Cell Signaling Technology | 3035 | 1:1000 |
| NF-κB2 p100/p52 | Cell Signaling Technology | 4882 | 1:1000 |
| p-NF-κB2 p100 | Cell Signaling Technology | 4810 | 1:1000 |
| NIK | Cell Signaling Technology | 4994 | 1:1000 |
| p-IKKα/$\beta$ (Ser176/180) | Cell Signaling Technology | 2697 | 1:1000 |
| IKKα | Cell Signaling Technology | 11930 | 1:1000 |
| IKKβ | Cell Signaling Technology | 8943 | 1:1000 |
| p-IκBα (Ser32) | Cell Signaling Technology | 2859 | 1:1000 |
| IκBα | Cell Signaling Technology | 4814 | 1:1000 |
| TNF-R1 | Cell Signaling Technology | 3736 | 1:1000 |
| LTβR | Proteintech | 20331-1-AP | 1:1000 |

**Supplementary Table 3.**
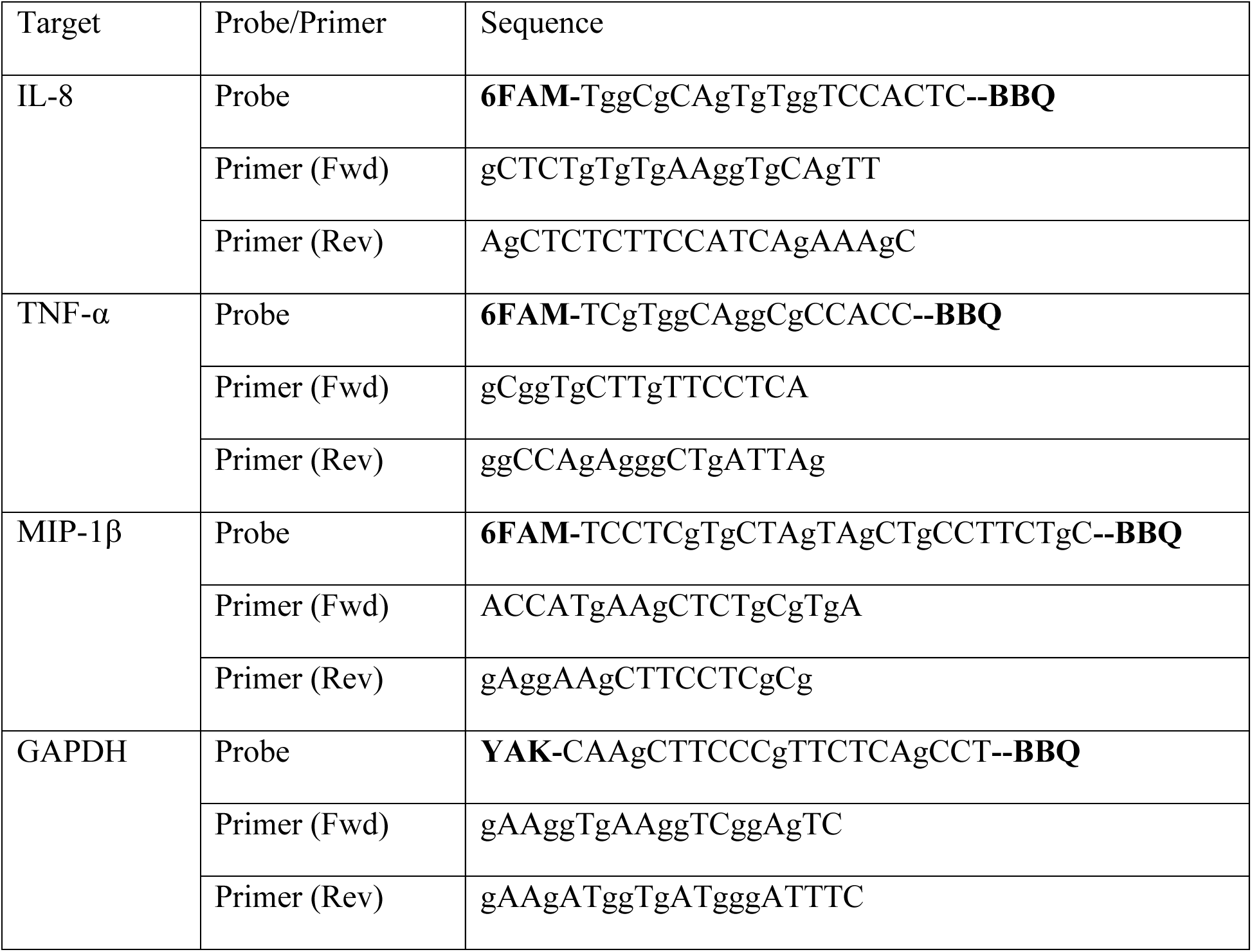
Sequences of TaqMan primers and primers.

